# StableProt: Structure-Aware Deep Learning for Protein Thermostability (*T_m_*) and Environmental Adaptation (OGT) Prediction

**DOI:** 10.64898/2026.11.06.764833

**Authors:** Bibhu Prasad Behera, Anshuman Dixit

**Affiliations:** Computational Biology and Bioinformatics Laboratory, iBRIC-Institute of Life Sciences (ILS), Bhubaneswar, 751023, Odisha, India

## Abstract

Predicting protein melting temperature (Tm) and organismal optimal growth temperature (OGT) is essential for enzyme engineering, biocatalyst design, and metagenomic bioprospecting, yet computational prediction faces several fundamental bottlenecks: sequence-only representations miss tertiary packing; dual-task coupling induces destructive gradient conflict; natural database skew (>87% mesophiles) drives mesophilic collapse; and existing tools produce uncalibrated point predictions. To address these challenges, we present StableProt, a structure-aware framework that couples amino acid sequences with Foldseek 3Di structural tokens via a frozen SaProt backbone to capture 3D contacts. Disjoint heads condition thermodynamic stability on ecological context without gradient conflict, while targeted 14% mesophilic subsampling expands thermophile representation to eliminate mesophilic collapse. Calibrated ensemble modeling reduces calibration error to 1.92%, providing dependable 95% confidence intervals. Evaluating models across 10°C temperature intervals reveals severe baseline collapse at thermal extremes, whereas StableProt maintains robust accuracy across regimes. On ProThermDB and zero-shot FireProtDB, StableProt achieves state-of-the-art probabilistic accuracy (CRPS = 4.52°C and 8.71°C) while recovering 92.0% of hyperthermophiles (≥80°C), and in prospective validation, its calibrated intervals capture both enzyme stability (75.3% coverage) and host environmental growth temperatures (78.1% coverage). StableProt is freely accessible as an interactive web server at https://stableprot.dixitlab.org/.

**Graphical Abstract:** 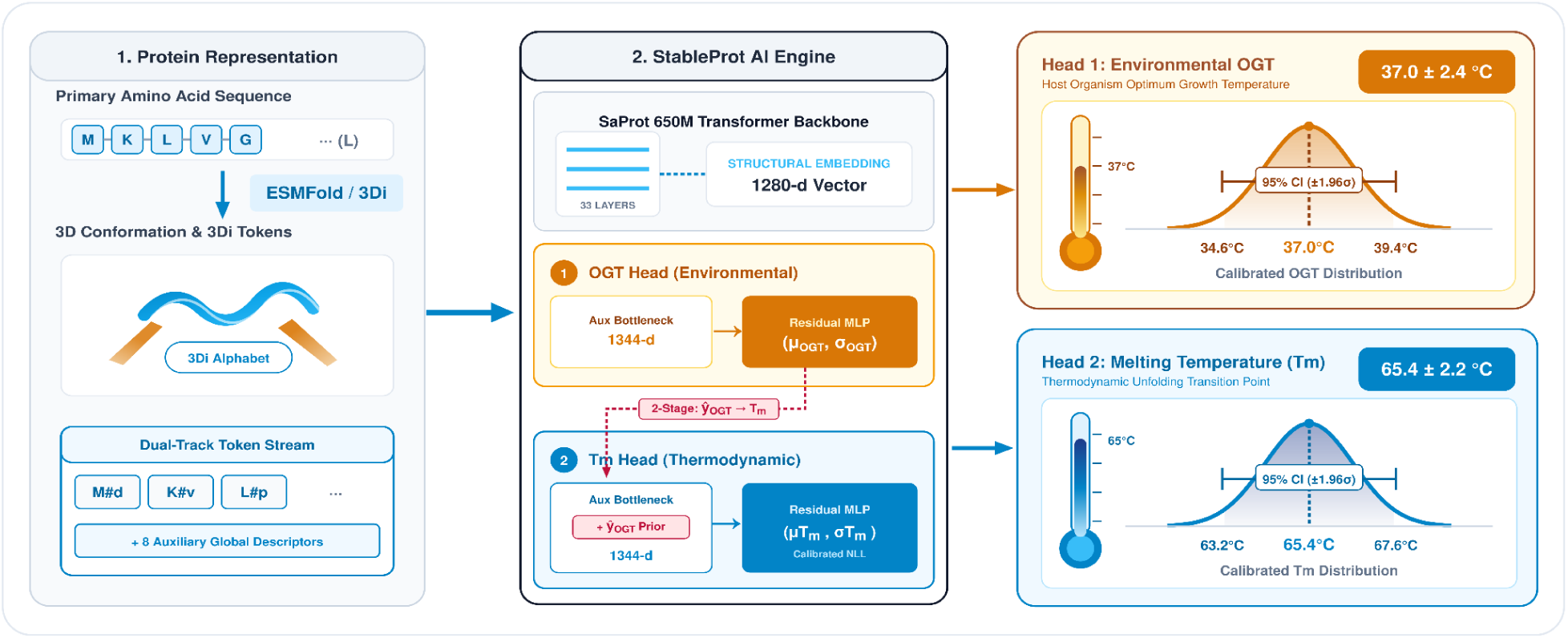

## Introduction

Thermal stability is a fundamental determinant of macromolecular structural integrity and functional persistence under high temperature (1, 2). Prediction of thermal stability is essential for the rational design and optimization of biomolecules for industrial, diagnostic, and therapeutic applications (3, 4). The melting temperature (T_m_) is a key quantitative descriptor, defined as the exact temperature at which 50% of a molecular population is unfolded at equilibrium (5). This macroscopic property is governed by intricate interactions, such as intramolecular hydrogen bonding, salt bridges, hydrophobic core packing, protein-protein interactions (PPIs), post-translational modifications (PTMs), bound ligands/cofactors and solvent conditions (pH, ionic strengths)(6–8). Optimal growth temperature (OGT), by contrast, is an ecological boundary of the host organism, defined as the temperature at which the organism exhibits maximum growth rate (9, 10). OGT is determined experimentally either by temperature-gradient incubation or by growth-rate profiling, in which growth rates are fitted to a temperature-response curve (10). Several methods, including Differential Scanning Fluorimetry (DSF), Circular Dichroism (CD) spectroscopy, and Differential Scanning Calorimetry (DSC), are labor-intensive. Recent methods such as mass spectrometry-based Thermal Proteome Profiling (TPP/ MS-CETSA) enable proteome-wide thermal stability profiling directly from cellular lysates (11–13). However, these approaches are costly, require high-end mass spectrometers, and rely on indirect curve fitting that is difficult to standardize to absolute T_m_ values (14). As a result, there are fewer than 50,000 curated experimental T_m_ entries across all public databases globally. By comparison, OGT-linked protein sequences number in the hundreds of millions, roughly three orders of magnitude more. Making effective use of this data is therefore essential for accurate computational prediction of thermal stability from sequence alone, where structural coverage falls short (15).

Early T_m_ prediction approaches relied on rule-based heuristics and statistical methods utilizing biophysical descriptors such as relative solvent accessibility (RSA), aliphatic indices, dipeptide composition, average hydrophobicity, predicted secondary structure fractions, and charge distribution to predict T_m_ values, yet yielded modest results(16, 17). With early algorithms having plateaued, and machine learning showing promise in other fields, researchers adapted these methods for biological tasks applying Decision Trees, Random Forests, Support

Vector Machines, gradient boosting methods such as XGBoost and LightGBM, and regularized regression on physicochemical feature sets (18, 19). However, these approaches were limited by hand-crafted feature engineering and lacked hierarchical, contextual, or evolutionary representations from sequences. Deep learning methods soon followed, applying convolutional, recurrent, and Transformer-based architectures directly to sequence data (20, 21). Yet, these approaches yielded only modest improvements without a fundamental breakthrough. That breakthrough came with protein language models (pLMs), whose usefulness emerged only with scale (22). These self-supervised Transformer models, analogous to Large Language Models (LLMs), are trained on massive protein sequence datasets capturing evolutionary syntax, conservation constraints, and residue co-evolutionary patterns. Key foundation pLMs such as ProtTrans(23), the ESM family(24), and ProtBERT(25) made training on small datasets viable, spawning specialized predictors such as TemStaPro(26), TemBERTure(14), DeepSTABp(27), ESMStabP(28), and ThermoFormer(29) (Transformer regressor for OGT/T_m_) that have shown great promise. More recently, the adoption of accurate structure prediction tools like AlphaFold(30) and ESMFold(24) has enabled structure-aware pLMs such as SaProt(31), which unifies amino acid sequences with the 20-state Foldseek 3Di structural alphabet into a dual-track discrete token representation. This structure-aware representation captures tertiary packing and non-local contacts, outperforming sequence-only models on structure-sensitive benchmarks.

OGT and T_m_ represent distinct domains with limited correlation(32). OGT supplies broad environmental context across thousands of organisms, yet this abundance is offset by noise. For instance, a bacterium with an OGT of 37°C routinely has proteins that remain active at 75-80°C, revealing the disconnect between organismal growth temperature and protein stability(33). The noise is compounded by how the data is sourced: OGT values are often reported at coarse resolution, derived from species-level taxons, with over 87% of records clustering between 25°C and 40°C, with 42.3% concentrated at exactly 28°C, 30°C, or 37°C. Crucially, because all proteins within an organism share the same OGT label, models relying on taxonomic metadata exhibit near-zero rank-order discrimination within a single species. In contrast, T_m_ is empirically measured and cleaner than OGT(34), yet it remains sparse. While it provides direct thermodynamic endpoints, its scarcity leaves most of the proteome uncharted. Consequently, models trained on OGT alone cannot resolve single-protein variations, while models trained on T_m_ alone lack out-of-distribution generalization. Neither dataset alone is adequate, yet each compensates for the other’s weakness. OGT’s abundance offsets T_m_’s scarcity, and T_m_’s precision corrects OGT’s coarseness. However, naively merging them in a single regression task introduces physical domain distortion, as they represent distinct physical quantities with only modest correlation. StableProt overcomes this conflict by deploying parameter-isolated disjoint heads that preserve their distinct physical identities, while allowing ecological context to anchor thermodynamic predictions without backward gradient interference.

Beyond representing distinct domains, OGT data is heavily skewed toward the mesophilic band (35). Under a standard symmetrical loss, models minimize error by predicting values close to the mesophilic mean. We term the resulting failure mode mesophilic collapse: the systematic compression of the predicted dynamic range toward the dataset’s mesophilic prior mean, induced by class imbalance under a symmetrical loss function. Models suffering from this collapse fail to generalize to thermal extremes, yet mask their degradation behind deceptively low overall MAE scores. These limitations necessitate a modelling strategy that respects the distinct nature of each label while leveraging their complementarity.

Following the advent of pLMs, the initial predictor adopted multi-bracket classification, using threshold classifiers across the thermal stability spectrum(26). While this model reported favorable AUC, it showed limited generalization when evaluated independently. Binned threshold predictions offer no continuous stability estimate, limiting their utility for enzyme engineering and industrial screening. Subsequent models moved to regression, enabling continuous T_m_ or OGT prediction(14, 27–29, 36). These models achieved improved predictive performance across diverse proteins. However, despite these advances, none of these approaches provide uncertainty bounds, nor are they statistically calibrated. Compounding these issues, benchmark evaluations often suffer from sequence leakage between training and test sets. Under rigorous sequence separation (<30% identity), predictive accuracy across these models drops substantially, particularly outside the mesophilic mode. This degradation is masked by the whole-dataset MAE, which is dominated by the mesophilic majority, and only emerges under temperature-binned evaluation that exposes model behaviour across thermal regimes. At the architectural level, joint optimization on a shared backbone forces conflicting gradient updates, as their gradients point in opposing directions (cos*θ* < 0), and optimizing one objective actively degrades the representational fidelity of the other. Overcoming this conflict requires an architecture that preserves their distinct physical identities while allowing them to inform each other.

To address these challenges, we developed **StableProt**, the first dual-head predictor to provide statistically calibrated confidence intervals for both T_m_ and OGT prediction **(Figure 1)**. StableProt resolves the limitations with four strategies. First, StableProt leverages SaProt’s Foldseek 3Di tokens to capture the 3D folding geometry and non-local packing essential to thermal denaturation that 1D sequence models miss. Second, to prevent overfitting and gradient conflict, we keep the 650M-parameter SaProt backbone frozen and deploy disjoint neural heads with zero shared weights (cos*θ* = 0) **(Figure 1)**. During inference, the predicted OGT is fed as an auxiliary prior link into the T_m_ head, allowing ecological context to anchor thermodynamic predictions without backward gradient interference (37). Third, to eliminate mesophilic collapse, we apply biophysical consistency filtering and subsample mesophilic data to 14%, expanding thermophilic representation by 2.3-fold (16.8% to 38.0%) and balancing learning across thermal extremes **(Figure 2)**. Fourth, StableProt couples a heteroscedastic variance module and a five-seed ensemble to capture both aleatoric and epistemic uncertainty, with confidence intervals calibrated via cross-fitted scaling **(Figure 5)**. We evaluated StableProt using the Continuous Ranked Probability Score (CRPS), which assesses distributional calibration and evaluates deterministic point predictors at their exact MAE (CRPS = MAE) with zero mathematical penalty. Across in-domain holdouts (ProThermDB)(38) and zero-shot out-of-distribution (OOD) benchmarks such as FireProtDB(39), FLIP Meltome(40, 41), BRENDA(42), and <30% identity partitions, StableProt achieves state-of-the-art accuracy and calibration, surpassing sequence-only and multi-task predictors **(Figure 3**, **Figure 4**, **Figure 6)**. In prospective validation, StableProt’s calibrated 95% confidence intervals achieved 75.3% coverage on industrial enzymes (61/81 scored variants; **Table 5A**) and 78.1% coverage on environmental markers (25/32; **Table 5B**), accurately predicting both T_m_ and OGT directly from sequence. StableProt includes an interactive web application suite **(Figure 7)** for *in silico* mutagenesis and stabilizing hypothesis generation before wet-lab synthesis.

**Figure 1.**
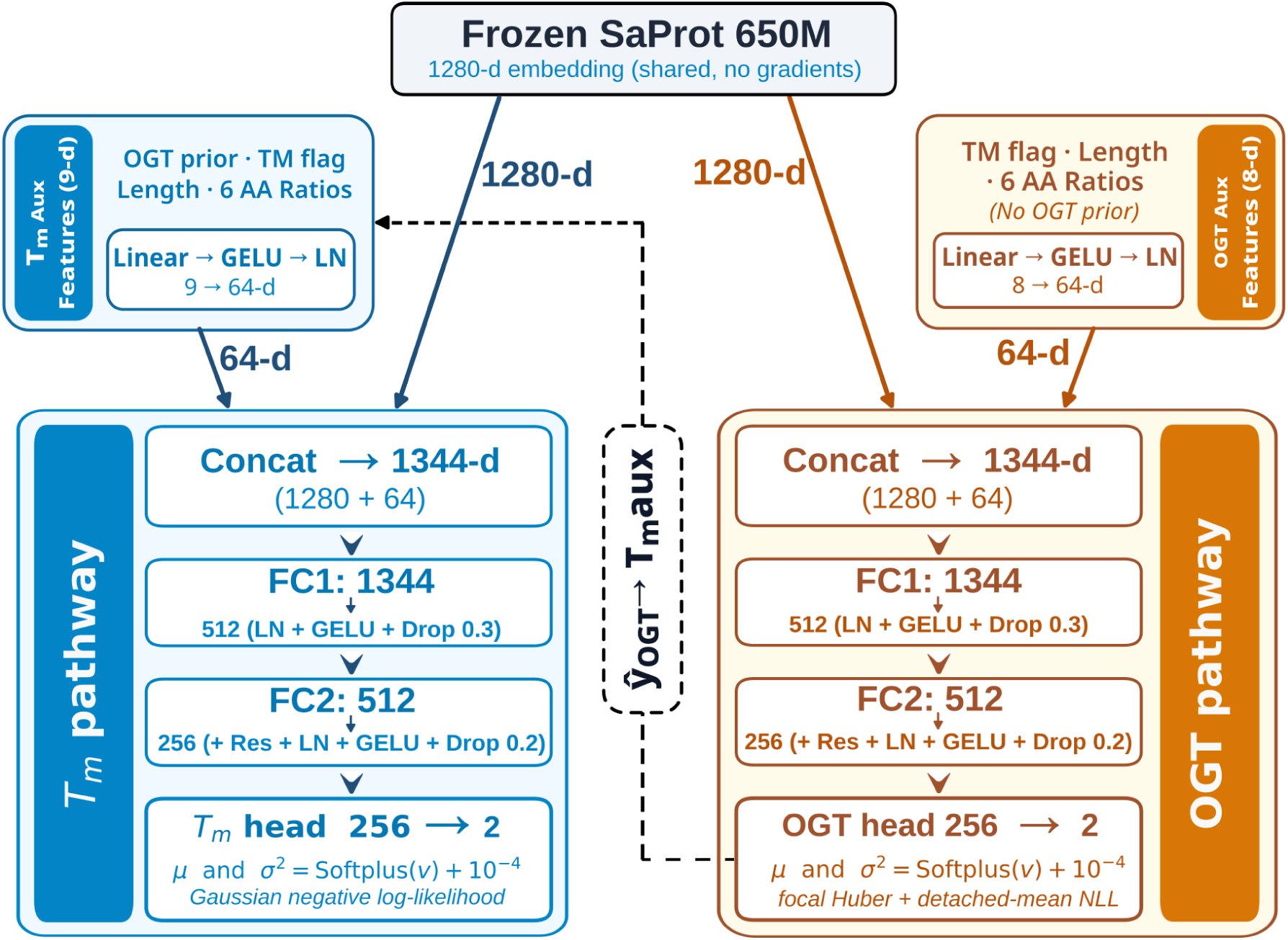
Standalone Disjoint Architecture of the StableProt Framework. A frozen SaProt 650M foundation backbone generates 1,280-d sequence embeddings. Auxiliary feature projections (9 → 64-d for T_m_, 8 → 64-d for OGT) yield 1,344-d latent vectors feeding independent three-layer residual MLP heads. The structure-conditioned T_m_ head predicts mean T̂_m_ and aleatoric variance *σ²*_Tm_ under Gaussian NLL. The sequence-only OGT head optimizes a Focal Huber loss with a detached-mean variance head. A unidirectional noise-corrupted link injects predicted OGT context into T_m_ as a prior. Five-seed ensembling and variance scaling produce calibrated 95% confidence intervals.

**Figure 2.**
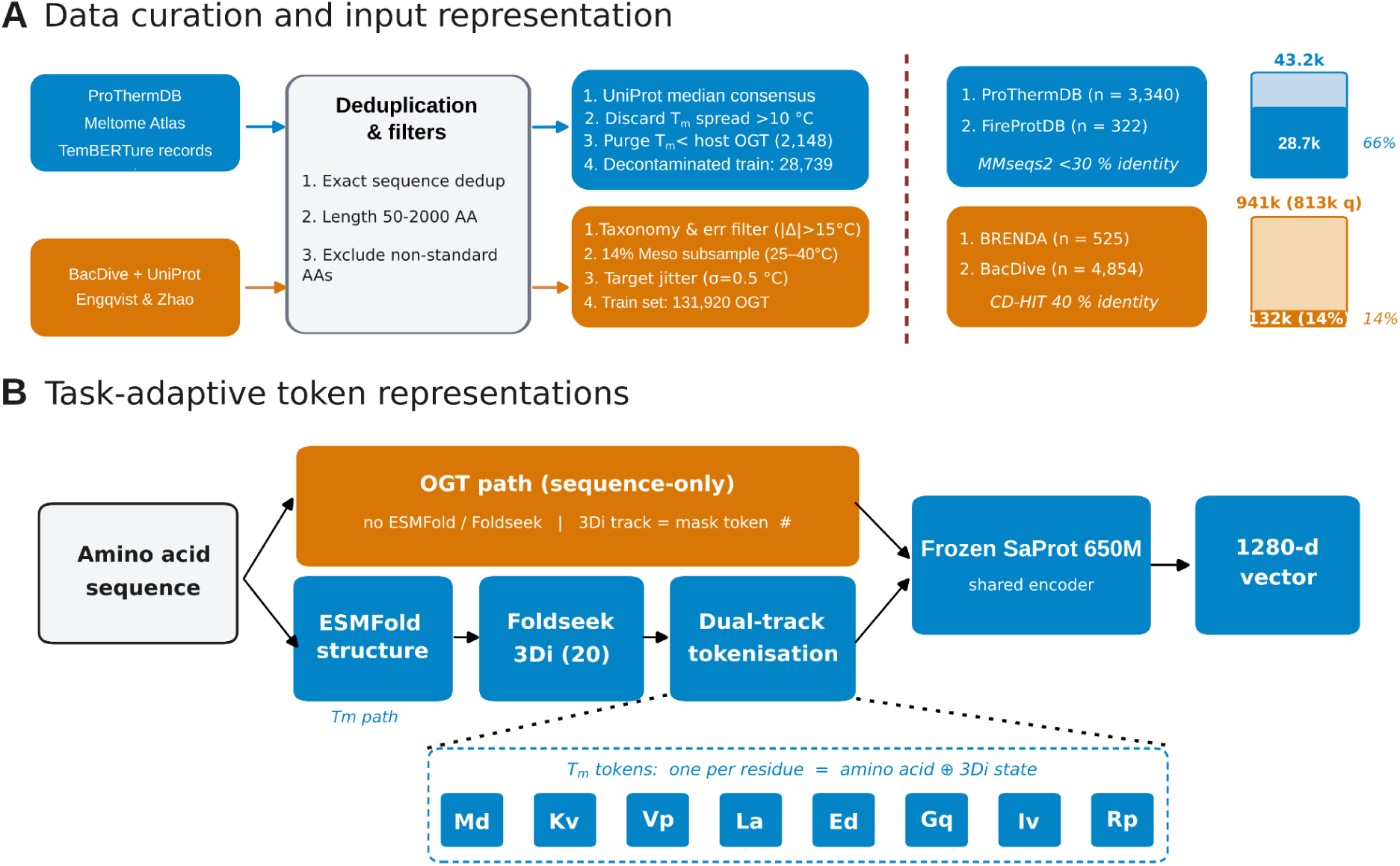
Targeted Data Curation Pipeline and Input Representation (A) Data curation and filtering: Multi-stage workflow across T_m_ and OGT databases with deduplication, length filtering (50-2,000 residues), replicate consensus, and biophysical sanitization (T_m_ ≥ OGT, purging 2,148 records). A 14% mesophilic subsampling rate (25-40°C) shifts thermophile batch representation from 16.8% to 38.0%. Bidirectional homology decontamination (MMseqs2 30%, CD-HIT 40%) against evaluation holdouts ensures zero data leakage. **(B) Task-adaptive representation:** The T_m_ head pairs amino acid sequences with Foldseek 3Di structural alphabet tokens ([Aa][3Di]), capturing 3D conformational packing. The OGT head uses sequence-level structural masking (aa#) without 3Di tokens. Both streams pass through the shared, frozen SaProt 650M backbone.

**Figure 3.**
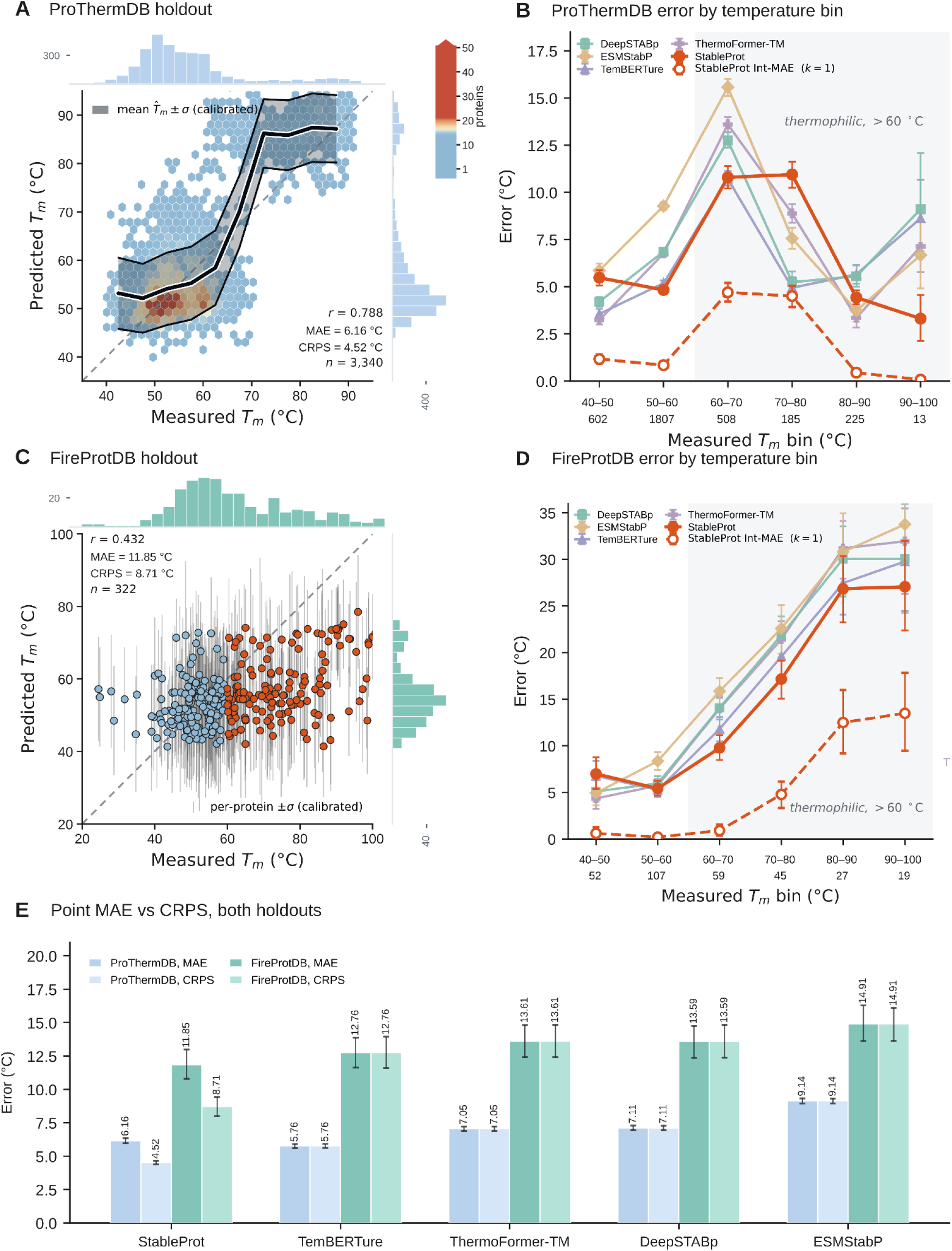
Unified Empirical Melting Temperature (T_m_) Benchmarks. **(A) ProThermDB holdout:** Parity density hexbin on 3,340 non-redundant proteins (MAE = 6.16°C, CRPS = 4.52°C, Pearson r = 0.788; OGT holdout r = 0.824) with calibrated binned error ribbon (*μ ± σ*) and marginal density histograms. **(B) ProThermDB binned error:** Mean Absolute Error across 10°C intervals (40-50°C to 90-100°C) benchmarking StableProt against TemBERTure, ThermoFormer-TM, DeepSTABp, and ESMStabP, highlighting flat error into the shaded thermophilic zone (>60°C) alongside confidence-adjusted Int-MAE (*k=1*). **(C) FireProtDB holdout:** Zero-shot out-of-distribution parity scatter on 322 proteins (MAE = 11.85°C, CRPS = 8.71°C) with per-protein calibrated uncertainty bars (±σ), marginal distributions, and discrete regime coloring (blue for <60°C, red for ≥60°C). **(D) FireProtDB binned error:** Per-bin MAE comparison across temperature intervals, demonstrating StableProt’s steady accuracy compared to the steep error escalation exhibited by competing regressors at elevated temperatures. **(E) Point MAE vs CRPS, both holdouts:** Paired bar chart across both evaluation holdouts (ProThermDB and FireProtDB). For deterministic point forecasts, CRPS equals point MAE; only StableProt emits calibrated confidence intervals, allowing CRPS to split below point MAE (4.52°C vs. 6.16°C on ProThermDB; 8.71°C vs. 11.85°C on FireProtDB).

**Figure 4.**
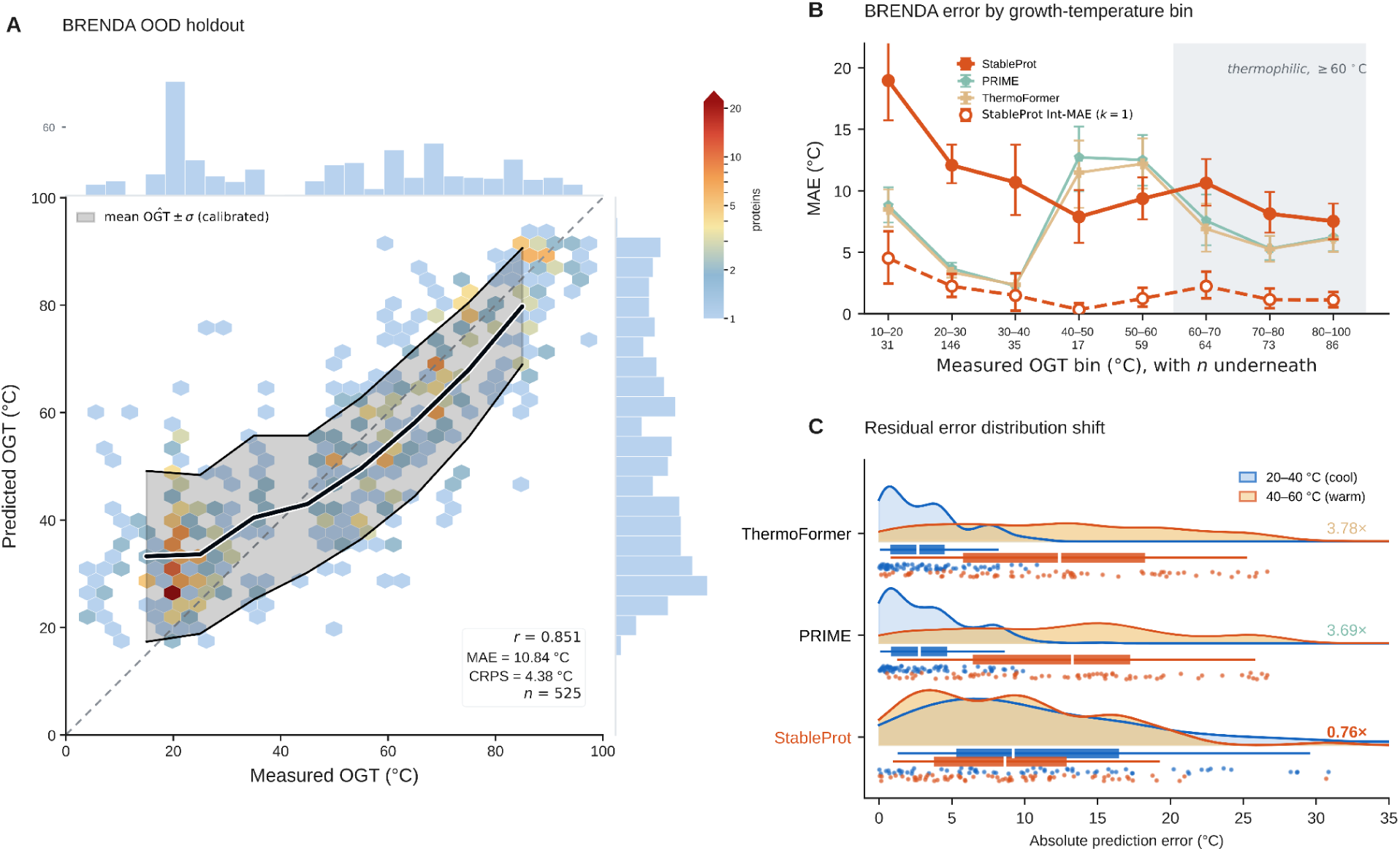
Organismal Growth Temperature Generalization and Collapse Analysis. **(A) BRENDA OOD holdout:** Measured vs. predicted OGT scatter parity on the external enzyme benchmark (N = 525, overall MAE = 10.84°C, bin-wise CRPS = 8.23°C, r = 0.851). **(B) Error by growth-temperature bin:** Binned MAE across eight 10°C intervals, highlighting that competing models surge nearly fourfold above 40°C while StableProt maintains a flat error profile. **(C) Residual error distribution shift:** Error distributions and thermal collapse ratio [MAE(40-60°C) / MAE(20-40°C)] contrasting mesophilic (20-40°C) against warm (40-60°C) organisms (0.76✕ for StableProt vs. 3.78✕ for ThermoFormer and 3.69✕ for PRIME).

**Figure 5.**
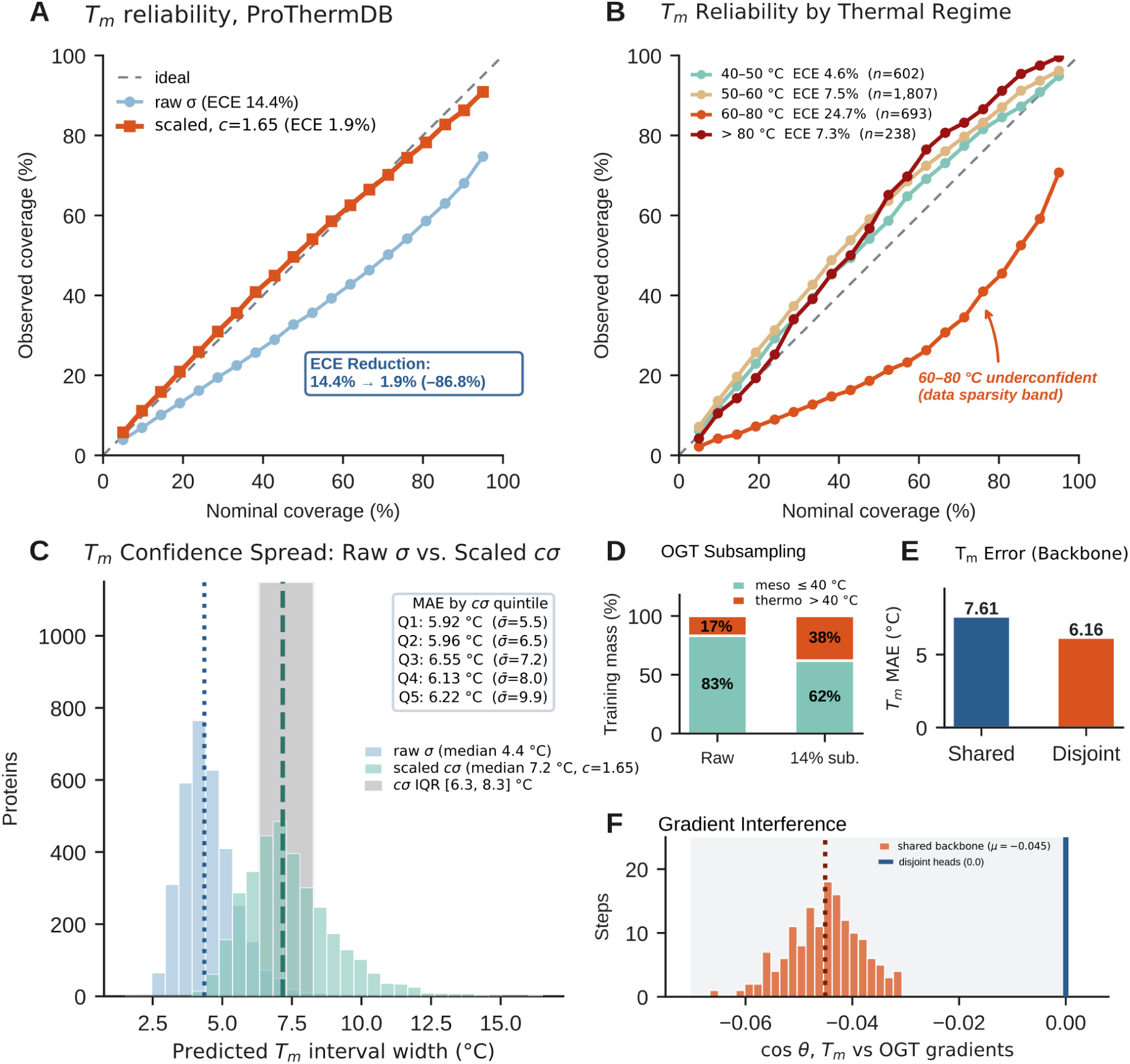
Uncertainty Calibration, Reliability, and Architectural Ablations. **(A) Reliability diagram:** Nominal vs. observed coverage on ProThermDB. Variance scaling (*c\** = 1.65) reduces Expected Calibration Error from 14.4% to 1.92% (86.8% reduction). **(B) Reliability by thermal regime:** Stratified coverage across stability bands, demonstrating reliable intervals with slight conservatism in the 60-80°C band. **(C) Confidence spread:** Calibrated uncertainty distribution, showing confidence intervals widen monotonically across error quintiles. **(D) Mesophilic subsampling ablation:** The 14% subsampling shifts thermophile batch representation from 16.8% to 38.0%, accelerating convergence by 55% and reducing hyperthermophile error (> 80°C) by 1.83°C. **(E) Shared-backbone ablation:** Jointly optimizing T_m_ and OGT on a single backbone incurs a +1.50°C penalty on ProThermDB (6.11°C → 7.61°C, single-seed run), eliminated by disjoint MLP heads. **(F) Gradient interference:** Histogram of task loss gradient cosine similarities during shared-backbone training (cos*θ^-^* = −0.077), confirming destructive gradient conflict in disjoint heads (cos*θ^-^*≡ 0).

**Figure 6.**
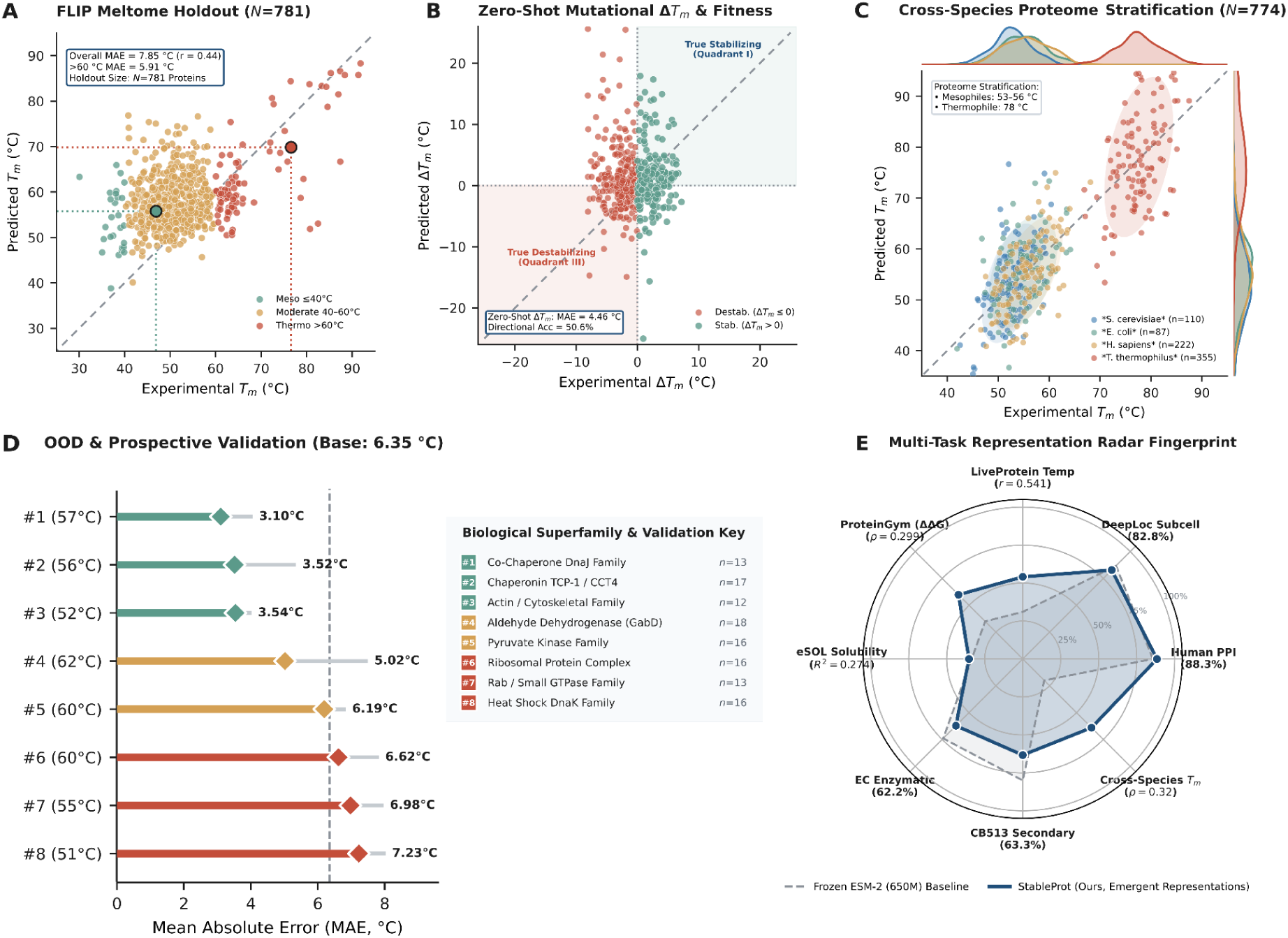
Zero-Shot Out-of-Distribution Transfer, Proteome Parity, and Mutation Sensitivity. **(A) FLIP Meltome holdout (N = 781):** Cellular thermal shift assay screen showing experimental vs. predicted T_m_, with error dropping to 5.91°C on thermophiles (>60°C; overall MAE 7.85°C, r = 0.44). **(B) Zero-shot mutational ΔT_m_ and fitness:** Single-point mutation dataset on FireProtDB (N = 3,649; MAE 5.05°C, 56.8% directional accuracy). Inset: ProteinGym **ΔΔ**G rank correlation (ρ = 0.299 StableProt vs. 0.175 ESM-2). **(C) Cross-species proteome parity (N = 774):** Scatter plot with 68% contour ellipses and marginal KDEs across four model organisms (*S. cerevisiae, E. coli, H. sapiens, T. thermophilus*), confirming intra-species variation resolution. **(D) Performance-sorted OOD validation (Base: 6.35°C):** Plot of biological superfamily key covering eight MMseqs2 fold clusters sorted by error tier. **(E) Multi-task performance radar chart:** Polar chart benchmarking frozen SaProt representations against ESM-2 (650M) across eight downstream functional and biophysical tasks.

**Figure 7.**
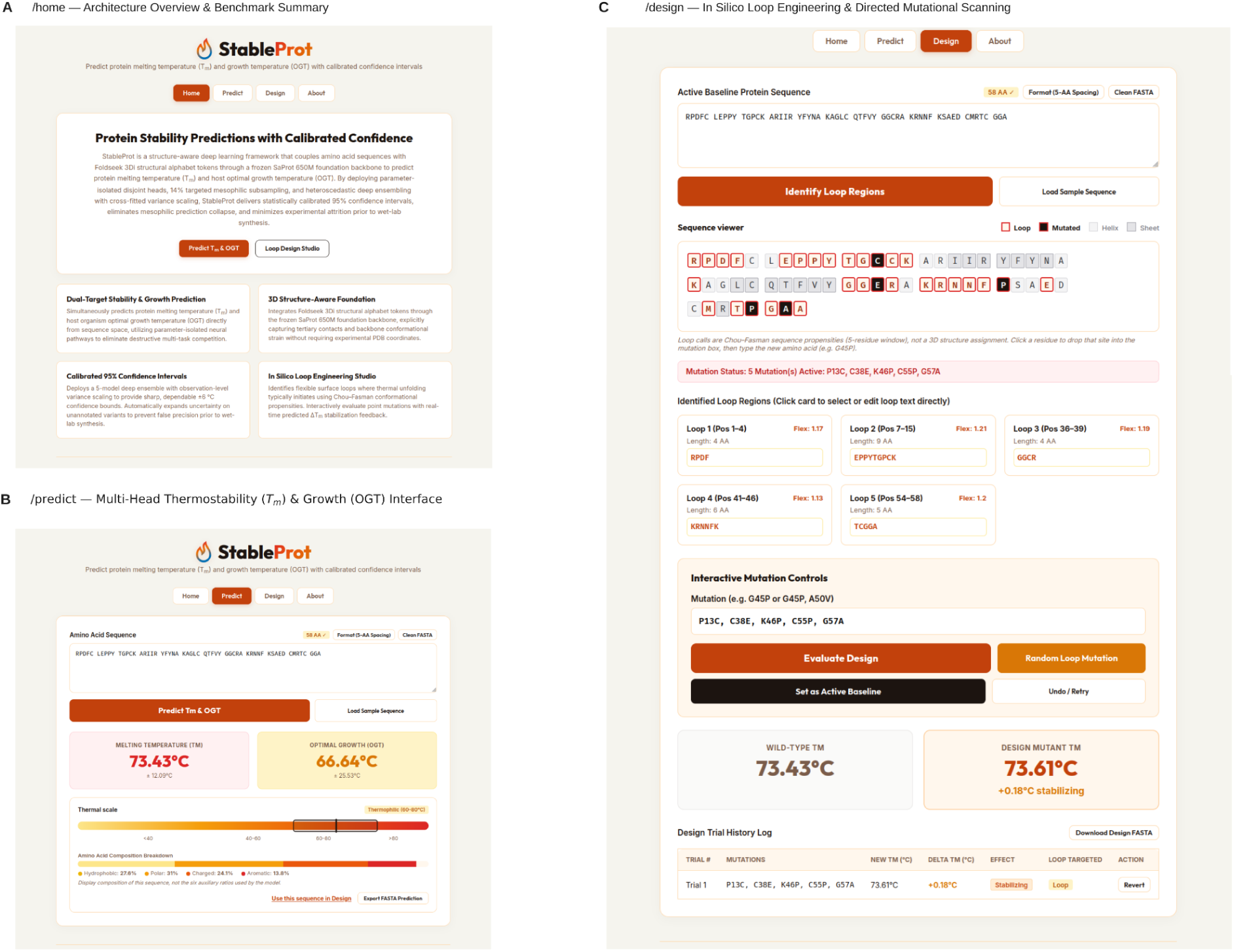
Containerized Interactive Web Application Suite. **(A) /home** : Architecture & Benchmarks: System overview interface presenting the dual-head architecture, frozen SaProt backbone, disjoint heads, and performance summaries across test cohorts. **(B) /predict** : Multi-Head Inference Interface: Two-stage prediction portal providing automated sequence sanitization, structure-aware inference, and calibrated 95% confidence intervals for T_m_ and OGT. **(C) /design** : *In Silico* Loop Engineering Toolkit: Protein engineering workspace integrating Chou-Fasman secondary structure propensity scoring, flexible loop detection, and single-point mutation evaluations.

## Materials and Methods

### Data source and curation

For this study, we assembled two datasets for each T_m_ and OGT respectively by curating measurements from different repositories. Sequences across both these datasets were cleaned of any non-canonical amino acid characters (X, B, Z, J), and lengths were constrained to 50-2,000 residues to align with the modeling capacity of the language model. For proteins appearing in both T_m_ and OGT datasets, records were co-partitioned into training and evaluation splits to prevent cross-task leakage **(Table S2)**. T_m_ data was compiled from ProThermDB (38), Meltome Atlas (40), and literature-curated records from TemBERTure (14), totaling 43,229 raw unfolding measurements. Records that shared identical UniProt accessions or identical primary sequences were clustered. Each consensus cluster T_m_ was established via the sample median, and clusters with >10°C internal spread were discarded as noisy outliers. We further purged sequences for which T_m_ < OGT, eliminating 2,148 biophysically inconsistent records **(Figure 2A; Supplementary Figure S1)**. Bidirectional homology decontamination resulted in a final set of 28,739 leak-free training sequences **(Table S2)**. For benchmarking, 3,340 ProThermDB entries were held out for in-domain testing, and an independent set of 322 sequences from FireProtDB was curated for out-of-distribution (OOD) evaluation. We evaluated the T_m_ head against four continuous regression models (TemBERTure, ThermoFormer-TM, DeepSTABp, and ESMStabP) as well as the multi-threshold classifier TemStaPro **(Table 1)**. Models were restricted to models with working code and weights; other tools lacked repositories or failed to execute.

**Table 1:** Architectural and methodological comparison of evaluated protein thermostability and growth temperature predictors. This table benchmarks StableProt against six state-of-the-art models across key architectural dimensions, including pre-trained backbone, parameter scale, input modality, and multi-task coupling. StableProt is the only framework that incorporates Foldseek 3Di structural tokens to explicitly capture tertiary folding cooperativity alongside primary sequences. Furthermore, unlike deterministic point predictors, it deploys parameter-isolated dual MLP heads that emit continuous Gaussian distributions with statistically calibrated 95% confidence intervals.

| Metric | TemStaPro | DeepSTABp | TemBERTure | ESMStabP | ThermoFormer | PRIME | StableProt (Ours) |
|---|---|---|---|---|---|---|---|
| **Publication Year** | 2024 | 2023 | 2024 | 2025 | 2026 | 2023 | 2026 (this work) |
| **Pre-trained Backbone** | ProtT5-XL | ESM-1b | ProtBERT | ESM-2 650M | Custom Transformer | ESM-1b | SaProt 650M |
| **Backbone Parameters** | 3.0 B | 650 M | 420 M | 650 M | 96 M | 690 M | 650 M |
| **Input Modality** | 1D Sequence | Sequence + Lysate TPP | 1D Sequence | 1D Sequence | 1D Sequence | Sequence + Taxonomy | Sequence+Structure (3Di) |
| **3D Structural Features** | No | No | No | No | No | No | Yes (Foldseek 3Di) |
| **Prediction Target** | *T<sub>m</sub>* threshold class | *T<sub>m</sub>* | *T<sub>m</sub>* | *T<sub>m</sub>* | OGT & *T<sub>m</sub>* (separate ckpts) | OGT | Both *T<sub>m</sub>* and OGT |
| **Multi-Task Coupling** | Single-task | Shared parameters | Single-task | Single-task | Shared parameters | Single-task | Disjoint per-task MLP heads |
| **Output Representation** | 6 binary brackets | Point forecast (ŷ) | Point forecast (ŷ) | Point forecast (ŷ) | Point forecast (ŷ) | Point forecast (ŷ) | Continuous distribution (*μ*, *σ<sup>2</sup>*) |
| **Uncertainty Quantification** | None | None | None | None | None | None | Calibrated 95% interval (±1.96*c*\*σ) |
| **Evaluated Checkpoint** | Standard release | DeepSTABp_v1 | TemBERTure | ESMStabP | GinnM/ThermoFormer-TM | AI4Protein/Prime_690M | v9_disjoint (5-seed ensemble) |

OGT data was curated from BacDive (43), NCBI taxonomy metadata, and published microbial growth temperature dataset from Engqvist (44), totaling 1.4 million raw records. Strains deviating >20°C from their taxonomic family median OGT were discarded as likely measurement error, which would otherwise inject noise and impair model learning. After length and residue filtering, the corpus contained 941,041 records. However, this corpus is heavily skewed: 42.3% of records sit at exactly 28, 30, or 37°C, and 87% fall between 25 and 40°C. Standard training on this distribution induces mesophilic collapse. To prevent this, we applied a targeted mesophilic subsampling rate of 14% during training. Each epoch randomly samples 14% of mesophilic sequences (25-40°C) while retaining 100% of psychrophilic (<25°C) and thermophilic (>40°C) sequences. This shifts thermophile representation from 16.8% to 38.0%, producing 131,920 records per epoch (**Figure 2A**). Finally, because roughly 84% of database entries are rounded integers, continuous Gaussian target jitter (σ = 0.5°C) was added during training to smooth discrete integer rounding artifacts, and targets were Z-score normalized. From this curated corpus, 5% (N = 47,052) was reserved as an internal validation split, and 525 enzyme sequences across 263 organisms from BRENDA were curated as an external out-of-distribution test set. The OGT head was evaluated against PRIME(36) and ThermoFormer(29)**(Table 1**, **Table 3)**.

To evaluate model performance on enzyme engineering targets, we designed an independent validation suite of 82 non-redundant sequences across four diverse enzyme cohorts: high-activity marine carrageenases(45, 46), literature dataset containing marine carrageenases(47, 48), and thermostable and thermolabile lipases curated from the Lipase Engineering Database(49) **(Table 5A and Supplementary Data 1)**. In parallel, to validate environmental thermal adaptation, we evaluated 32 cellular biomarker sequences spanning Adenylate Kinase (50), Ribosomal protein uL2 (51), HSP60 and GroEL chaperonins (52) **(Table 5B)**.

### Bidirectional Homology Decontamination

To ensure evaluation reflects true generalization rather than memorization, training and evaluation partitions were separated by bidirectional homology decontamination **(Figure 2B)**. Because sequence similarity artificially inflates benchmark metrics, we enforced strict identity thresholds at the limits of detectable homology. For T_m_ data, we applied MMseqs2(53) at ≤30% identity (coverage = 0.8) to eliminate structural fold memorization within the twilight zone. Conversely, for the 1.4-million OGT corpus, we used CD-HIT(54) at ≤40% identity as per the tool’s algorithmic clustering limit and to avoid over-purging taxonomic diversity.

Accordingly, our T_m_ evaluation benchmarks comprising the held-out ProThermDB (N = 3,340) and zero-shot FireProtDB (N = 322) were screened against both the T_m_ training pool and the entire OGT corpus at ≤30% identity. This cross-task boundary prevents information leakage during two-stage inference, ensuring the upstream OGT head cannot pass memorized representations to the downstream T_m_ head. Finally, all competing predictors (TemStaPro, TemBERTure, DeepSTABp, ESMStabP, and ThermoFormer) were re-evaluated on these exact decontaminated holdouts, ensuring that comparative metrics reflect true architectural capacity rather than split discrepancies.

## Structure-Aware 3Di Tokenization

We next addressed the limitations of sequence-only representations. Thermal denaturation is governed by tertiary structure. As sequence-only language models lack structural coordinates, their stability predictions are constrained by 1D sequence representations. Therefore, we extracted 1,280-dimensional representations using the frozen SaProt 650M backbone. SaProt uses a dual-track token alphabet that combines the 20 amino acid characters with 20 discrete structural states derived from Foldseek 3Di discretization. SaProt performs best when both sequence and structural tokens are provided, but it remains functional with sequence tokens alone.

Structural 3Di tokens were generated exclusively for the T_m_ dataset, as folding the 1.4-million OGT corpus was computationally prohibitive. Structures were predicted primarily using ESMFold(24), using OpenFold (55) for long sequences exceeding GPU memory limits. Coordinates were then discretized into 3Di tokens using Foldseek. To align with SaProt’s standard 1,024-token context window, sequences exceeding this length were trimmed (<0.4%), with Foldseek’s mask token (#) assigned to any missing coordinates. (**Figure 2B**).

### Model architecture

StableProt decouples T_m_ and OGT prediction into two disjoint regression heads sharing zero trainable weights(56)(**Figure 1**). This isolation eliminates gradient interference, as the shared-trunk control exhibited conflicting updates (cos*θ* = -0.077) and a +1.50°C MAE penalty on T_m_ **(Figure 5E, 5F)**. This also enables **one-way information flow**: the OGT head runs first, passing its prediction into the T_m_ head as a physical prior without gradient interference. Each head uses a lightweight independent multi-layer perceptron (MLP)(57) with the SaProt backbone kept frozen. To support SaProt representations with task-specific constraints, auxiliary feature vectors were added with Z-score normalization. The OGT head receives **eight scalars**: a binary transmembrane topology flag, sequence length, and six composition fractions (charged, hydrophobic, proline, cysteine, aromatic, and normalized Aliphatic Index), where the Ikai aliphatic index is divided by 100 to rescale values to [0, 1]:

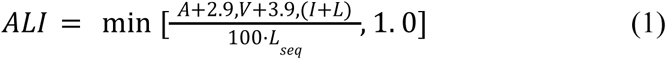

where *A, V, I*, and *L* denote residue counts of alanine, valine, isoleucine, and leucine, respectively; *L_seq_* is the total sequence length; 2.9 and 3.9 represent the relative side-chain volumes of aliphatic residues compared to alanine; and min[⋅,1.0] bounds rare hyper-aliphatic outliers. The T_m_ head uses the same eight features plus the predicted OGT scalar, totaling nine dimensions. Each auxiliary vector is projected through a linear bottleneck (mapping 9 to 64 for T_m_ and 8 to 64 for OGT), followed by LayerNorm and GELU. This projection expands the auxiliary features to 64-dim, preventing them from being overwhelmed when concatenated with the 1,280-dim mean-pooled SaProt embedding to form a 1,344-dim input. Each head then processes this input through a three-layer residual MLP that maps the 1,344-dimensional input through hidden layers of 512 and 256 dimensions to the output. Each residual block applies a skip linear projection from 1,344 to 256 dimensions, followed by LayerNorm; dropout of 0.3 and 0.2 is applied after the first and second layers, respectively. Linear layers were initialized using Kaiming normal initialization (58), which scales weights by fan-in to preserve gradient variance through GELU activations. A 3-layer configuration achieved the lowest validation loss **(Table S1).** Each head outputs a variance estimate alongside its mean prediction, used to quantify the aleatoric and epistemic uncertainty as formalized below.

### Loss Functions & Bounded Variance

T_m_ and OGT have distinct data distributions and noise profiles. Each head is therefore trained with a specialized loss objective. The T_m_ head outputs a point prediction *μ* and an unconstrained raw log-variance *ν*, which prevents variance collapse. It is optimized using a heteroscedastic Gaussian negative log-likelihood loss:

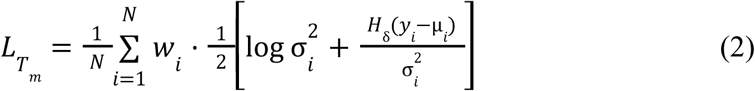

where *N* is the batch size, *y_i_* is the experimental melting temperature, *μ_i_* is the predicted mean, σ_i_^2^ is the predicted aleatoric variance, and w_i_ is the sample reliability weight. Here, H_δ_(r_i_) is the Huber residual on error *r_i_=y_i_−μ*_i_ [defined as 1/2*r ^2^* for ∣*r_i_*∣≤*δ,* and *δ*(∣*r_i_*∣−1/2*δ*) for ∣*r_i_*∣>*δ*], with threshold *δ*=5°C set above the assay variance (1-3°C) to preserve quadratic sensitivity for small errors while limiting outlier gradients, and 1/2log*σ* ^2^ penalizes arbitrary variance inflation. Next, we applied an inverse interquartile range (IQR) sample weight w_i_:

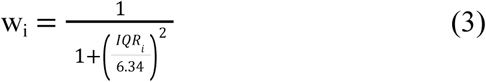

where IQR*_i_* is the interquartile range of reported experimental T_m_ values across replicate literature measurements for protein *i*, and *w_i_*∊(0,1] scales sample reliability. The scale constant 6.34°C corresponds to the 98.5th percentile of IQR across proteins with multiple measurements (15,405 sequences), dampening conflicting records. For proteins with only a single measurement, we imputed the dataset mean IQR of 0.62°C (yielding w_i_≈0.99). IQR weighting was only applied to T_m_, as OGT is an organism-level annotation without per-protein replicates. The OGT head is trained using a Focal Huber loss (59) in Z-score space, incorporating a transition threshold of *δ* = 15°C, a focusing factor *γ* = 2.0, and a threshold *β* = 0.5, where temperature bins are weighted by a frequency multiplier:

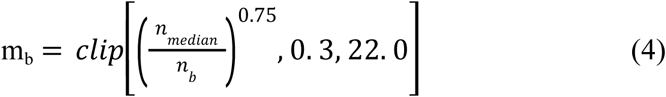

where n_b_ is the sample count in 10°C temperature bin *b*, nmedian is the median bin count across the training set, the 0.75 power provides sublinear frequency damping, and clip[⋅,0.3,22.0] bounds multipliers between 0.3✕ and 22.0✕. This provides up to 22-fold gradient amplification for rare hyperthermophiles (>80°C) while preventing gradient instability. To enable variance prediction for OGT without degrading regression accuracy, we combined the Focal Huber loss with a detached Gaussian NLL(60) variance objective:

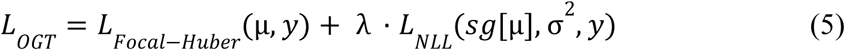

where L_focal-Huber_ is the weighted regression loss with focusing exponent *γ*=2.0, L_NLL_ is the Gaussian negative log-likelihood 1/2[log*σ*^2^+(*y*−sg[*μ*])^2^/σ^2^], *λ*=1.0 balances the two objectives, and sg[⋅] denotes the stop-gradient operator (∂/∂*θ*sg*[μ]*=0). The stop-gradient was applied throughout training, because under mesophilic imbalance the variance head inflated *σ²* to reduce large residuals, degrading point accuracy (MAE worsened from 13.6°C to 21.4°C). Detaching *μ* from the variance gradient preserves point accuracy while still producing calibrated uncertainty. Variance was bounded identically to T_m_, with initial value *σ²* ≈ 0.693.

### Training and optimization

All models were implemented in PyTorch and trained on an NVIDIA RTX 6000 Ada GPU. Each head was optimized independently using AdamW(61) with a base learning rate of 1×10^-4^, weight decay of 1×10^-5^, and batch size of 64. Learning rates followed a cosine annealing schedule with warm restarts (T_0_ = 10, T_mult_ = 2, η_min_ = 1×10^-6^), and training was terminated when validation loss failed to improve for 15 consecutive epochs **(Supplementary Figure S2)**. Gradients were clipped to a maximum norm of 1.0. During training, input embeddings were regularized with Gaussian noise (*σ* = 0.02, p = 0.15), and target labels were smoothed with additive jitter (*σ* = 0.5°C). Architectural and optimization hyperparameters are catalogued in **Table S1**.

During inference, the OGT head runs first and supplies its standardized prediction to the T_m_ auxiliary vector. Throughout training, the T_m_ head receives the ground-truth OGT corrupted with additive Gaussian noise (*σ* = 2°C) to prevent the network from developing an over-reliance on a noise-free scalar prior and compounding prediction error during two-stage inference.

All predictions are generated by an ensemble of five independently trained models initialized with fixed random seeds (1–5). The ensemble prediction is the mean across models, and total predictive variance follows the Law of Total Variance, combining aleatoric and epistemic uncertainty. We report this observation-level predictive variance rather than the standard error of the ensemble mean, which would compress intervals by √5 ≈ 2.24✕ and cause nominal 95% intervals to cover only ∼62% of measurements.

### Model evaluation metrics and calibration

Model performance was evaluated using deterministic regression metrics such as Mean Absolute Error (MAE), Root Mean Squared Error (RMSE), Pearson correlation coefficient (r), Spearman rank correlation (ρ). For candidate screening, we evaluated MCC, Top-10% screening precision, and screening F1 (at T_m_ ≥ 80°C). Statistical significance was assessed via paired bootstrap resampling with 95% percentile intervals. Since mesophiles dominate the dataset, we report both overall and bin-wise metrics (averaged across 10°C intervals) to assess accuracy across thermal extremes **(Table 3**, **Table 4)**. To compare StableProt’s predictive distributions against deterministic models on a shared scale, we used the Continuous Ranked Probability Score (CRPS), which measures the integrated squared error between the predicted cumulative distribution and the empirical step function. For a Gaussian forecast:

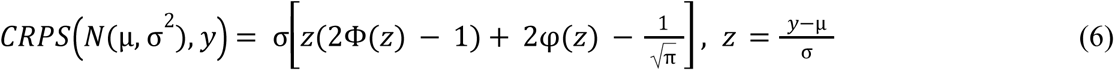

where *y* is the experimental target, *μ* and *σ* are the predicted mean and standard deviation, *z* is the normalized prediction residual, Φ(z) and φ(z) represent the standard normal cumulative distribution and probability density functions, respectively, and 1/√π accounts for the expected pairwise dispersion between independent draws. Two properties make CRPS suitable for cross-model benchmarking. First, as σ → 0, CRPS converges to *|y - μ|*, meaning deterministic models receive their MAE without penalty. Second, CRPS is minimized only when the predicted distribution matches the data distribution, so models cannot inflate variance to improve their score. To assess sharpness alongside error, we compute an interval-adjusted MAE at *k=1*, which penalizes only errors beyond the ±1σ predictive interval. Because networks trained under likelihood objectives often produce miscalibrated uncertainty, we fit a per-benchmark variance scale factor *c\** via two-fold cross-fitting and report Expected Calibration Error (ECE) across 20 nominal confidence levels. Empirical scaling factors (ProThermDB = 1.65; FireProtDB = 3.62) are detailed in **Supplementary Table S4** and **Supplementary Figure S3**, yielding calibrated 95% confidence intervals.

## Results

### Evaluation on melting temperature benchmarks

We benchmarked StableProt against five competing models, comparing four regressors (TemBERTure, ThermoFormer-TM, DeepSTABp, and ESMStabP) and one classifier (TemStaPro) (**Table 1**). All models were evaluated on the decontaminated ProThermDB holdout (N = 3,340; **Figure 3A**), with TemStaPro assessed separately in the binary classification screening analysis **(Supplementary Table S3).** StableProt achieved a CRPS of 4.52°C (MAE = 6.16°C, Pearson r = 0.788; **Table 2**). Because competing models produce deterministic outputs without uncertainty estimates, their CRPS scores are identical to their MAE values: TemBERTure (5.76°C), ThermoFormer-TM (7.05°C), DeepSTABp (7.11°C), and ESMStabP (9.14°C) **(Table 2)**. StableProt’s CRPS of 4.52°C represents a 1.24°C reduction over the nearest competitor (TemBERTure) and a 2.53°C-4.62°C improvement over the remaining models, indicating that modeling aleatoric variance effectively accounts for experimental assay noise. Across 10°C intervals, StableProt maintained a balanced error profile into elevated regimes, recording the lowest error in extreme thermophiles (**Figure 3B**). On rank-order consistency, StableProt achieved a Spearman correlation of ρ = 0.517, matching TemBERTure (ρ = 0.516) and exceeding DeepSTABp (ρ = 0.500) and ESMStabP (ρ = 0.402) **(Table 2)**.

**Table 2.** Accuracy and probabilistic benchmarking across in-domain and zero-shot out-of-distribution *T_m_* holdouts. Models were evaluated on in-domain ProThermDB (n = 3,340) and zero-shot out-of-distribution FireProtDB (n = 322) across continuous error metrics and thermophilic screening criteria. StableProt achieves the lowest Continuous Ranked Probability Score on both benchmarks (CRPS = 4.52°C and 8.71°C), substantially outperforming deterministic models whose CRPS collapses to their MAE. On zero-shot OOD targets, StableProt demonstrates superior structural generalizability, leading in MAE (11.85°C), hyperthermophile recall (81.8%), and industrial screening F1 (0.474).

| **Model** | **MAE (°C) ↓** | **CRPS (°C) ↓** | **Pearson r ↑** | **Spearman ρ ↑** | **Recall (≥80°C) ↑** | **Top-10% Prec ↑** | **Screening F1 ↑** |
|---|---|---|---|---|---|---|---|
| **ProThermDB, in distribution (n = 3,340)** |  |  |  |  |  |  |  |
| TemBERTure | **5.76** | 5.76 | 0.826 | 0.516 | 57.1% | **0.521** | 0.568 |
| ThermoFormer-TM | 7.05 | 7.05 | **0.829** | **0.563** | 90.3% | 0.449 | 0.618 |
| DeepSTABp | 7.11 | 7.11 | 0.812 | 0.500 | 55.9% | 0.497 | 0.535 |
| ESMStabP | 9.14 | 9.14 | 0.715 | 0.402 | 86.1% | 0.512 | 0.617 |
| **StableProt (+ calibrated σ)** | 6.16 | **4.52** | 0.788 | 0.517 | **92.0%** | 0.494 | **0.650** |
| **FireProtDB, zero-shot out of distribution (n = 322)** |  |  |  |  |  |  |  |
| TemBERTure | 12.76 | 12.76 | 0.369 | 0.214 | 42.9% | 0.286 | 0.343 |
| DeepSTABp | 13.59 | 13.59 | **0.436** | 0.239 | 45.5% | 0.263 | 0.333 |
| ThermoFormer-TM | 13.61 | 13.61 | 0.393 | 0.233 | 63.6% | 0.250 | 0.359 |
| ESMStabP | 14.91 | 14.91 | 0.330 | 0.236 | 54.5% | 0.273 | 0.364 |
| **StableProt (+ calibrated σ)** | **11.85** | **8.71** | 0.432 | **0.350** | **81.8%** | **0.333** | **0.474** |

**Note:** Top-10% precision: among the 10% of sequences with the highest predicted T<sub>m</sub>, the fraction with true T<sub>m</sub> ≥ 80°C.

In enzyme engineering, identifying candidates at T_m_ ≥ 80°C is a primary goal(62). On this screening task, StableProt recovered 92.0% of hyperthermophiles on ProThermDB (recall = 0.920, F1 = 0.650), outperforming ThermoFormer-TM (recall = 90.3%, F1 = 0.618), ESMStabP (recall = 86.1%, F1 = 0.617), TemBERTure (recall = 57.1%, F1 = 0.568), and DeepSTABp (recall = 55.9%, F1 = 0.535) **(Table 2)**.

### Evaluation on organismal optimal growth temperatures

Growth temperature data is heavily skewed toward mesophiles, with over 87% of records falling between 25-40°C (**Figure 2A**). Therefore, overall MAE alone can mask mesophilic collapse, as models achieve low error by interpolating the mesophilic mode while losing accuracy at thermal extremes. To evaluate OGT prediction in this context, we assessed the OGT prediction head across three partitions: internal holdout (val_ogt_, N = 47,052), decontaminated BacDive benchmark (N = 4,854), and the external out-of-distribution BRENDA enzyme set (N = 525) (**Table 3**, **Table 4**, **Figure 4B**). To characterize this, we report both overall and bin-wise metrics averaged across 10°C intervals. The collapse pattern is unambiguous on the full holdout, PRIME achieved 2.1-2.4°C error within the mesophilic mode (71.1% of data) but increased to 8.7-21.6°C outside it (overall MAE = 4.75°C, bin-wise MAE = 10.08°C) **(Table 4)**. The same pattern appeared on BacDive (2.1-2.4°C inside the mesophilic band versus 9.8-22.9°C outside, bin-wise MAE = 10.38°C) and on BRENDA (2.26°C in the 30-40°C band versus 11.5-20.9°C outside, bin-wise MAE = 8.56°C) **(Table 4)**. The recurrence of this frequency-dependent error pattern across two architectures and three independent partitions is systematic rather than stochastic and constitutes a signature of **mesophilic collapse.**

**Table 3.** Overall and bin-wise-averaged performance across external and internal optimal growth temperature (OGT) benchmarks. Predictors are evaluated on zero-shot BRENDA (n = 525) and internal BacDive (n = 4,854) holdouts using both count-weighted (overall) and temperature-balanced (bin-wise) metrics. While deterministic models minimize overall MAE by collapsing toward the dense mesophilic majority (∼37°C), StableProt achieves the lowest bin-wise CRPS on both datasets (6.54°C on BacDive and 8.23°C on BRENDA). This demonstrates that targeted 14% mesophilic subsampling prevents mesophilic collapse, maintaining balanced predictive fidelity across the thermal spectrum.

| **Benchmark and model** | **MAE**<br>**overall↓** | **MAE**<br>**bin-wise↓** | **CRPS**<br>**overall↓** | **CRPS**<br>**bin-wise↓** | **r↑** | **ρ↑** | **scale c** |
|---|---|---|---|---|---|---|---|
| BRENDA, out of distribution (n = 525) |  |  |  |  |  |  |  |
| ThermoFormer | **6.48** | 8.29 | **6.48** | 8.29 | **0.938** | **0.934** | none |
| PRIME | 6.75 | 8.56 | 6.75 | 8.56 | 0.934 | 0.933 | none |
| StableProt | 10.84 | 11.79 | 7.55 | **8.23** | 0.851 | 0.835 | 1.22 |
| BacDive, internal test (n = 4,854) |  |  |  |  |  |  |  |
| ThermoFormer | **4.71** | 9.81 | **4.71** | 9.81 | - | - | none |
| PRIME | 4.96 | 10.38 | 4.96 | 10.38 | - | - | none |
| StableProt | 7.94 | **9.23** | 5.63 | **6.54** | - | - | 0.87 |

StableProt avoids this collapse through 14% mesophilic subsampling. On the full OGT holdout, StableProt achieved an overall MAE of 6.12°C (bin-wise MAE = 9.55°C; r = 0.824, ρ = 0.781), leading in 6 of 10 thermal regimes overall (and 7 of 10 against PRIME) **(Table 4)**. On BacDive, StableProt achieved a bin-wise MAE of 9.23°C, outperforming ThermoFormer (9.81°C) and PRIME (10.38°C) **(Table 3).** ThermoFormer and PRIME recorded lower overall MAE (4.71°C and 4.96°C, compared to 7.94°C for StableProt) by allocating capacity to the dense mesophilic center, but this apparent advantage vanished under bin-balanced evaluation. StableProt outperformed both models in seven of ten temperature intervals on BacDive, including all thermophilic regimes from 40°C to 100°C. **(Table 4**, **Figure 4B)**

**Table 4.** Granular per-temperature-bin error analysis across the thermal spectrum (0-100°C). Mean Absolute Error is resolved across ten 10°C intervals across three independent OGT cohorts (N = 47,052, N = 4,854, and N = 525). The breakdown reveals that model accuracy is driven almost entirely by the dense mesophilic window (20-40°C), while their error spikes dramatically across extremophilic regimes. In contrast, StableProt leads in 6 of 10 (Full Holdout) and 7 of 10 (BacDive Benchmark) thermal regimes, dominating 5 of 6 thermophilic and hyperthermophilic regimes (>40°C) via frequency-reweighted Focal Huber loss. † Full Holdout: Baseline overall MAE is driven by the 20-40°C mesophilic subset (71.1% of data; errors 2.11-2.36°C). Under unweighted bin-wise evaluation, StableProt achieves the lowest macro MAE (9.55°C vs. 9.62°C for ThermoFormer and 10.08°C for PRIME) and leads in 6 of 10 bins. Holdout RMSE = 8.74°C, r = 0.824, ρ = 0.781. ‡ BRENDA Benchmark: Baselines outperform StableProt across most temperature intervals on BRENDA (Macro MAE 8.29°C and 8.56°C vs. 11.79°C), as StableProt’s mesophilic subsampling causes systematic over-prediction in the 0-40°C range. StableProt leads only in the 40-60°C intervals (7.91°C and 9.41°C vs. 11.48-12.72°C) and achieves comparable bin-wise CRPS (8.23°C vs. 8.29°C and 8.56°C) via calibrated uncertainty bounds.

| **Temperature Bin** | **Sample Counts (n)<br>Holdout / BacDive / BRENDA** | **Full Holdout<br>ThermoFormer / PRIME / StableProt** | **BacDive Benchmark<br>ThermoFormer / PRIME / StableProt** | **BRENDA OOD<br>ThermoFormer / PRIME / StableProt** |
|---|---|---|---|---|
| **0-10°C** | 11 / 1 / 14 | 21.52 / 21.59 / **17.95** | 22.57 / 22.93 / **15.15** | 21.03 / **20.86** / 25.75 |
| **10-20°C** | 318 / 32 / 31 | **11.51** / 11.80 / 12.98 | **10.53** / 11.03 / 12.21 | **8.47** / 8.80 / 19.04 |
| **20-30°C** | 17,735 / 1,736 / 146 | **2.11 /** 2.13 / 7.27 | **2.05** / 2.09 / 7.79 | **3.38** / 3.68 / 12.17 |
| **30-40°C** | 15,580 / 1,675 / 35 | **2.28 /** 2.36 / 7.30 | **2.28** / 2.35 / 8.16 | 2.35 / **2.26** / 10.64 |
| **40-50°C** | 3,904 / 392 / 17 | 10.29 / 10.62 / **8.35** | 10.62 / 10.99 / **8.16** | 11.48 / 12.72 / **7.91** |
| **50-60°C** | 4,841 / 532 / 59 | 11.05 / 11.71 / **8.34** | 11.51 / 12.19 / **6.98** | 12.19 / 12.51 / **9.41** |
| **60-70°C** | 2,332 / 250 / 64 | 10.23 / 11.15 **/ 8.78** | 11.29 / 12.43 / **8.90** | **6.91** / 7.56 / 10.61 |
| **70-80°C** | 1,197 / 133 / 73 | **7.82** / 8.66 / 8.27 | 8.66 / 9.84 / **7.80** | **5.27** / 5.30 / 8.16 |
| **80-90°C** | 769 / 73 / 59 | 8.21 / 9.14 / **7.76** | 6.60 / 7.28 / **6.37** | **6.49** / 6.67 / 7.98 |
| **90-100°C** | 354 / 30 / 27 | 11.18 / 11.66 / **8.56** | 12.00 / 12.62 / **10.80** | 5.29 / **5.25** / 6.53 |
| **Summary Rows** |  |  |  |  |
| *Overall MAE* ↓ | 47,052 / 4,854 / 525 | **4.53** / 4.75 / 6.12† | **4.71** / 4.96 / 7.94 | **6.48** / 6.75 / 10.84‡ |
| *bin-wise MAE* ↓ | 10 bins | 9.62 / 10.08 / **9.55** | 9.81 / 10.38 / **9.23** | **8.29** / 8.56 / 11.79 |
| *CRPS bin-wise* ↓ | 10 bins | 9.62 / 10.08 / **6.81** | 9.81 / 10.38 / **6.54** | 8.29 / 8.56 / **8.23** |
| **Intervals Leading ↑** | 10 bins | 4/10 \| 0/10 \| **6/10** | 3/10 \| 0/10 \| **7/10** | 6/10 \| 2/10 \| **2/10** |

On the external BRENDA holdout (N = 525), StableProt underperforms both baselines on absolute error metrics, recording an overall MAE of 10.84°C and a bin-wise MAE of 11.79°C, compared to 6.48°C and 8.29°C for ThermoFormer, and 6.75°C and 8.56°C for PRIME **(Table 3**, **Table 4)**. ThermoFormer and PRIME also achieved substantially higher linear correlation (r = 0.938 and 0.934, vs. 0.851 for StableProt), benefiting from large-scale sequence pre-training that preserves broad phylogenetic rank ordering. StableProt’s higher error on BRENDA stems primarily from the 0-40°C intervals (43% of the cohort), where aggressive 14% mesophilic subsampling leads the model to systematically over-predict growth temperatures (averaging ∼34°C for 20-30°C enzymes; **Table 4**). StableProt leads only in the 40-60°C intervals, where baseline errors surge to 11.5-12.7°C while StableProt records 7.91°C (40-50°C) and 9.41°C (50-60°C). Across the full range, because StableProt outputs calibrated uncertainty bounds rather than deterministic point estimates, its bin-wise CRPS (8.23°C) remains comparable to the baselines (8.29°C for ThermoFormer, 8.56°C for PRIME).

For binary extremophile screening, StableProt reliably identified organisms adapted to elevated temperatures. On BacDive, screening for thermophiles (≥60°C) yielded an AUC of 0.894. On BRENDA, screening at ≥50°C achieved an AUC of 0.939 (PR-AUC = 0.950, F1 = 0.865), while screening at ≥60°C achieved an AUC of 0.946 (PR-AUC = 0.936, F1 = 0.828). These results demonstrate that mesophilic subsampling paired with frequency-reweighted Focal Huber loss prevents the OGT head from overfitting to the mesophilic majority, providing a robust screening filter for thermophilic biocatalysts.

### Uncertainty calibration and reliability analysis

We assessed calibration quality across nominal confidence levels (*p* ∈ [0.05, 0.95]) on both in-domain and out-of-distribution holdouts. Prior to calibration, raw network variances were substantially under-dispersed (overconfident), yielding an Expected Calibration Error (ECE) of 14.4% on ProThermDB. Applying two-fold cross-fitted variance scaling (*c\** = 1.65) reduced ECE to 1.92%, aligning empirical coverage with theoretical expectations **(Figure 5A)**: nominal 68% and 95% intervals enclosed 67.5% and 91.4% of experimental measurements, respectively (mean 95% half-width: 14.8°C; **Supplementary Figure S3**). Alternatively, global temperature scaling (*T* = 3.8) reduced benchmark-wide ECE to 0.46% (*R²* = 0.992; **Supplementary Table S4)**.

We next examined whether predicted uncertainty reflects informational ambiguity rather than uniform dispersion **(Figure 5B, 5C)**. Stratifying ProThermDB predictions by thermal stability revealed that predicted variance scaled with domain complexity **(Figure 5B)**, mesophilic targets (≤40°C) exhibited a standard deviation of σ = 5.21°C, whereas targets above 60°C exhibited an elevated mean standard deviation of σ = 7.14°C. Across uncertainty quintiles, prediction errors remained stable , indicating that the model produces wider confidence bounds when encountering targets with heterogeneous experimental records or atypical sequence composition **(Figure 5C)**.

### Ablation of architecture and training strategies

To quantify the contributions of our individual modeling choices, we systematically ablated the representation backbone, architectural connectivity, loss formulations, and training interventions **(Figure 5D-F; Table S1 and Table S4)**. Here, we evaluate the four primary mechanisms governing the model’s progression.

Structural conditioning delivered the largest single gain **(Table S4)**. Progressing from a baseline 1D sequence MLP (ProThermDB MAE = 12.28°C; FireProtDB MAE = 28.21°C) through intermediate refinements (LayerNorm, skip connections, and the 64-dimensional auxiliary bottleneck) reduced MAE to 6.84°C and 12.69°C, respectively. Incorporating SaProt 3Di structural tokens further decreased MAE to 6.11°C on ProThermDB and 10.84°C on FireProtDB, yielding a cumulative error reduction of 6.17°C in-domain and 17.37°C out-of-distribution.

Ablating the 14% mesophilic subsampling rate demonstrated its impact on thermal extremes **(Figure 5D)**. Restricting mesophile retention (25-40°C) to 14% per epoch shifted thermophile batch representation from 16.8% to 38.0%, accelerating loss convergence by 55% and reducing hyperthermophile prediction error (>80°C) from 9.83°C to 8.00°C, an improvement of 1.83°C.

Evaluating architectural connectivity confirmed the penalty of multi-task coupling **(Figure 5E)**. Jointly optimizing T_m_ and OGT on a single shared backbone degraded ProThermDB MAE by +1.50°C (6.11°C → 7.61°C) relative to independent training. Gradient tracking during joint optimization revealed an empirical cosine similarity of cos*θ* = -0.077 between the T_m_ Gaussian NLL gradient and the OGT Focal Huber gradient **(Figure 5F)**, confirming destructive gradient interference that parameter-isolated disjoint pathways eliminate.

Ablating the auxiliary OGT input confirmed the utility of environmental conditioning. Supplying the Tm head with true experimental OGT as an input feature yielded an MAE of 3.61°C, feeding the predicted OGT feature achieved 4.78°C, and substituting a constant mean feature (37.5°C) degraded accuracy to 5.00°C. In training, injecting Gaussian noise (σ = 2°C) into the OGT input yielded superior two-stage inference compared to training without noise corruption.

Additional ablations and hyperparameter evaluations are detailed in **Supplementary Table S1**, with remaining exploratory configurations provided in the accompanying codebase.

### Zero-shot out-of-distribution benchmarks

To evaluate generalization beyond public training distributions, we tested StableProt on two independent zero-shot benchmarks on the FireProtDB T_m_ holdout.

**FireProtDB** T_m_ **holdout (N = 322).** StableProt achieved MAE = 11.85°C and CRPS = 8.71°C (compared to 12.76-14.91°C across both metrics), while leading in rank-order correlation (ρ = 0.350 vs. 0.214-0.239) **(Table 2)**. Paired bootstrap testing over 4,000 resamples evaluated the per-protein error delta against each model, confirming that the 95% confidence interval of StableProt’s error reduction strictly excludes zero (*p* < 0.05; **Table 2)**.

Signed error distributions showed that competing models suffered from mesophilic collapse, systematically under-predicting thermophilic proteins **(Figure 3D)**. When transferring from in-distribution ProThermDB to out-of-distribution FireProtDB on thermophilic targets (70-80°C), prediction errors for competing models degraded severely: +14.63°C for TemBERTure, +16.62°C for DeepSTABp, and +15.02°C for ESMStabP **(Figure 3D)**. In contrast, StableProt limited transfer degradation to +5.45°C. Consequently, while competing sequence-based models achieved only 42.9% -63.6% recall on hyperthermophiles (≥80°C), StableProt maintained 81.8% recall **(Table 2)**.

### Generalization to external mutation and stability screens

We further evaluated StableProt across three independent external tasks: the FLIP Meltome high-throughput stability screen, homology-clustered family partitions, and physical single-point mutation stability shifts (ΔT_m_).

#### FLIP Meltome screen (N = 781)

On this proteome-scale cellular thermal shift assay benchmark, performance revealed clear limitations as overall correlation was modest (Pearson r = 0.44, Spearman ρ = 0.24, overall MAE = 7.85°C; **Figure 6A)**, and error was poor on cold-adapted proteins below 40°C (MAE = 16.02°C, n = 27), reflecting the severe scarcity of psychrophilic training data. However, accuracy improved on thermophiles above 60°C (MAE = 5.91°C, n = 94; 7.79°C between 40-60°C, n = 660), indicating that prediction fidelity does not degrade at high temperatures.

#### Homology cluster analysis

Partitioning the evaluation corpus into 5,861 disjoint homology clusters (≤30% sequence identity) yielded an overall MAE of 6.35°C, with steady accuracy across major structural fold families **(Figure 6D)**. Ranking enzymes within a single organism is harder, because every protein in a cell carries the same host OGT label and the same taxonomic background. Competing predictors returned negative intra-species rank correlations, and StableProt was the only model to stay positive in every tested species (ρ = 0.03-0.19; per-species MAE 6.55°C-10.36°C; **Figure 6C**). The 3Di tokens therefore appear to encode intrinsic fold stability rather than a host growth temperature prior.

#### Mutation ΔT_m_ benchmark

On the FireProtDB single-point mutation benchmark comprising 3,649 physical wet-lab-measured ΔT_m_ values from circular dichroism, differential scanning calorimetry, and fluorescence assays, performance highlighted an inherent architectural limitation. Computing independent wild-type and mutant predictions [Δ*T*_m_ = T_m_(mut) − T_m_(wt)] yielded an MAE of 5.05°C (RMSE = 7.78°C) and a directional sign accuracy of only 56.8% (AUC = 0.585, Pearson r = 0.19, Spearman ρ = 0.21; **Figure 6B**) marginally above random chance. This limitation arises because the network is trained on denaturation endpoints across diverse sequences rather than on mutational shifts, making it unsuitable for atomic-resolution **ΔΔ**G prediction. Instead, its wide uncertainty bounds serve primarily as a coarse triage filter. On the zero-shot ProteinGym substitution suite (**Figure 6B**), StableProt achieved a Spearman rank correlation of ρ = 0.299, outperforming the sequence-only foundation model ESM-2 650M (ρ = 0.175) and demonstrating that structure-aware representations better capture mutational fitness effects.

#### Multi-task representation probing

Finally, to test whether StableProt generalises to different general biological domains, we evaluated representations across a probing suite spanning protein-protein interactions (HumanPPI), cellular localization (DeepLoc), solubility (eSOL), enzyme classification (EC), secondary structure (CB513), fold topology (SCOP), experimental assay temperature tracking (LiveProteinBench), and zero-shot mutational fitness (ProteinGym) **(Figure 6E)**. The highest scores were obtained on interface and mutational fitness tasks: 88.3% on HumanPPI, ρ = 0.299 on ProteinGym, and r = 0.541 for assay temperature tracking on LiveProteinBench. Performance was lower on biophysical property prediction, with 82.8% on DeepLoc, *R²* = 0.290 on eSOL, and 64.6% residue accuracy on CB513. High-level evolutionary categorization yielded the lowest scores, at 63.8% on EC and 15.8% on SCOP. Relative to ESM-2 650M, StableProt achieved higher scores on ProteinGym and HumanPPI, comparable scores on DeepLoc and eSOL, and lower scores on CB513, EC, and SCOP. Taken together, these results indicate that stability-specific training retained general structural and functional information rather than degrading it.

### Experimental validation on enzyme cohorts

In addition to the benchmark dataset, we evaluated StableProt on 82 non-redundant experimental protein sequences spanning four diverse cohorts (82 experimental records; CgiB_Ce and CgkS appear in both the high-activity and from literature) **(Table 5A)**. These sequences represent real laboratory targets outside public training databases: two high-activity marine carrageenases (45, 46), thirteen marine carrageenases from corresponding literature (47, 48), and 69 industrial lipases (37 thermostable, 32 thermolabile) (49). Among the 82 unique sequences (84 experimental records), 81 possess published experimental references (with CaKC16B evaluated as a prospective blind target). Full sequence-level predictions, uncertainty bounds, and primary accessions are catalogued in **Table 5A** and **Supplementary Data 1.**

**Table 5.**
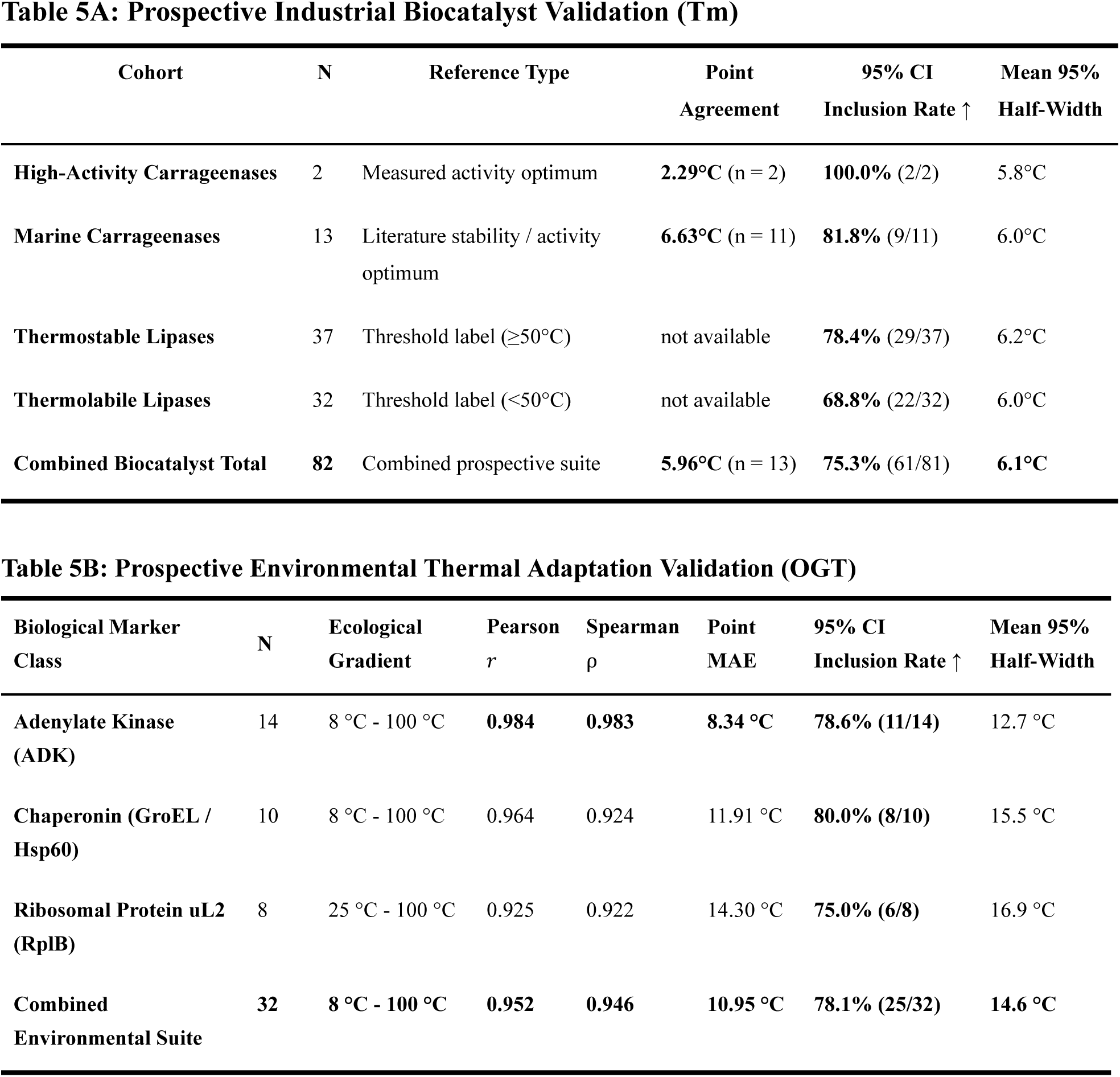
Prospective experimental validation across industrial biocatalysts (Table 5A) and environmental thermal niches (Table 5B). Table 5A evaluates single-molecule thermal stability (Tm) across industrial enzyme targets, with intervals denoting calibrated 95% confidence bounds (c = 1.56, μ ± 1.96·c·σ). Table 5B evaluates macroscopic environmental growth temperature (OGT) across global ecological thermal gradients (8 °C to 100 °C) inferred from single housekeeping biomarker sequences.

| **Cohort** | **N** | **Reference Type** | **Point Agreement** | **95% CI Inclusion Rate ↑** | **Mean 95% Half-Width** |  |  |
|---|---|---|---|---|---|---|---|
| **High-Activity Carrageenases** | 2 | Measured activity optimum | **2.29°C** (n = 2) | **100.0%** (2/2) | 5.8°C |  |  |
| **Marine Carrageenases** | 13 | Literature stability / activity optimum | **6.63°C** (n = 11) | **81.8%** (9/11) | 6.0°C |  |  |
| **Thermostable Lipases** | 37 | Threshold label (≥50°C) | not available | **78.4%** (29/37) | 6.2°C |  |  |
| **Thermolabile Lipases** | 32 | Threshold label (<50°C) | not available | **68.8%** (22/32) | 6.0°C |  |  |
| **Combined Biocatalyst Total** | **82** | Combined prospective suite | **5.96°C** (n = 13) | **75.3%** (61/81) | **6.1°C** |  |  |
| **Biological Marker Class** | **N** | **Ecological Gradient** | **Pearson *r*** | **Spearman ρ** | **Point MAE** | **95% CI Inclusion Rate ↑** | **Mean 95% Half-Width** |
| **Adenylate Kinase (ADK)** | 14 | 8 °C - 100 °C | **0.984** | **0.983** | **8.34 °C** | **78.6% (11/14)** | 12.7 °C |
| **Chaperonin (GroEL / Hsp60)** | 10 | 8 °C - 100 °C | 0.964 | 0.924 | 11.91 °C | **80.0% (8/10)** | 15.5 °C |
| **Ribosomal Protein uL2 (RplB)** | 8 | 25 °C - 100 °C | 0.925 | 0.922 | 14.30 °C | **75.0% (6/8)** | 16.9 °C |
| **Combined Environmental Suite** | **32** | **8 °C - 100 °C** | **0.952** | **0.946** | **10.95 °C** | **78.1% (25/32)** | **14.6 °C** |

On high-activity carrageenases with quantitatively characterized activity optima (N = 2), StableProt achieved an MAE of 2.29°C, with 100% inclusion within the calibrated 95% confidence intervals (2/2) **(Table 5A)**. For CgkS, the model predicted T_m_ = 47.38°C ± 6.33°C, within 2.38°C of its measured activity optimum (T_opt_ = 45°C). For CgiB_Ce, it predicted 42.19°C ± 5.36°C, within 2.19°C of T_opt_ = 40°C. We note that the catalytic activity optimum and thermal melting temperature are related but distinct physical endpoints, with T_opt_ typically falling below T_m_; this close correspondence reflects operational consistency rather than exact thermodynamic unfolding identity.

On the literature-curated marine carrageenase cohort (N = 13, 11 scored), StableProt achieved a continuous point MAE of 5.96°C across the 11 scored variants against parsed literature activity and stability ranges, with 81.8% of predictions (9/11) falling within the calibrated 95% confidence intervals. Two sequences in this cohort lacked reported literature stability ranges (CaKC16B, which is uncharacterized, and CgiB_Ce, evaluated against its catalytic optimum above) and were retained as prospective blind predictions. Combining all 13 carrageenases with continuous activity optima or stability ranges yielded an overall continuous MAE of 5.96°C (n = 13; **Table 5A**).

Evaluating uncharacterized industrial lipases highlighted the practical value of calibrated uncertainty bounds over rigid thresholding. For thermostable lipases (N = 37) and thermolabile lipases (N = 32), single point predictions clustered between 42°C and 49°C because candidate sequences lacked organismal growth annotations, causing the auxiliary OGT prior to default toward the mesophilic training mean (37-45°C). In such unannotated regimes, forced binary categorization across an arbitrary 50°C boundary yielded point classification accuracies of 43.2% (16/37) and 46.9% (15/32), respectively. In contrast, StableProt’s calibrated 95% confidence intervals expanded appropriately (±6.0°C to ±6.2°C half-width), successfully enclosing 78.4% of thermostable lipases (29/37) and 68.8% of thermolabile lipases (22/32) **(Table 5A, Supplementary Data 1)**. Rather than emitting false precision on ambiguous targets, the calibrated interval explicitly signals when candidate enzymes cannot be definitively resolved without empirical verification.

### Single-Gene Environmental Adaptation Profiling Across Global Ecological Niches

To test whether StableProt can infer an organism’s macroscopic environmental growth niche directly from single sequences, we profiled diverse housekeeping proteins: Adenylate Kinase (ADK) (50), Ribosomal protein uL2 (RplB) (51), HSP60 and Chaperonin GroEL (52) across model organisms spanning a wide temperature range (8°C to 100°C; **Table 5B**), with reference growth temperatures verified against BacDive (43) and Engqvist microbiological datasets (44). Across diverse habitats including Siberian permafrost (*Psychrobacter*, 15°C), temperate soil (*B. subtilis*, 37°C), desert soda lakes (*Natranaerobius*, 53°C), and abyssal hydrothermal vents (*Pyrococcus*, 100°C) StableProt demonstrated high rank-order fidelity from single ADK sequences alone (Pearson r = 0.984, Spearman ρ = 0.983, MAE = 8.34°C, 78.6% CI coverage [11/14]). On GroEL chaperonins, 80.0% of predictions (8/10) fell within calibrated 95% confidence intervals, and Ribosomal uL2 achieved 75.0% CI inclusion (6/8; r = 0.925).

Across the combined 32-marker environmental suite, StableProt achieved an overall Pearson r of 0.952, Spearman ρ of 0.946, and 78.1% 95% CI coverage (25/32). Predicted T_m_ values for mesophilic housekeeping proteins remained 10-20°C above predicted OGT (e.g., *E. coli* OGT = 37.4°C vs. Tm = 52.3°C), confirming that parameter isolation preserves the distinction between single-molecule denaturation and host macroscopic growth boundaries.

### Computational efficiency and web deployment

A major bottleneck limiting the adoption of deep learning tools in routine protein engineering is computational overhead. Many contemporary models require full parameter fine-tuning of multi-billion parameter foundation backbones or rely on uncurated pre-training on tens of millions of sequences. StableProt addresses this through parameter isolation and targeted data curation **(Table 1)**. By keeping the underlying SaProt 650M backbone completely frozen, the trainable footprint is restricted to the lightweight MLP projection heads. Each head contains approximately 1.17 million parameters, yielding a total trainable parameter count of 2.34 million across the system. This corresponds to a trainable parameter overhead of only 0.36% of the 650M backbone. Training efficiency is further enhanced by targeted mesophilic subsampling. While models such as ThermoFormer required pre-training across 96 million uncurated protein sequences, StableProt achieves competitive zero-shot accuracy using less than 1 million curated records. On a single GPU workstation (NVIDIA RTX 6000 Ada Generation), training the T_m_ head requires approximately 20 minutes per seed, roughly 1.5 hours for the complete five-seed ensemble, while the subsampled 941,041-record OGT head converges in approximately 2.5 hours per seed. In contrast, full fine-tuning of competing architectures such as ESMStabP (650 million parameters) or TemStaPro (3.0 billion parameters) requires substantial computational resources.

To facilitate widespread academic and industrial adoption, we containerized the complete StableProt inference engine into an open-source, production-ready web application (stableprot_deployment/, **Figure 7**). Built on a FastAPI backend with PyTorch runtime, the server is packaged with Docker and Docker Compose configurations supporting automated CUDA GPU acceleration with automatic CPU fallback. The interactive web interface provides a streamlined workflow for biological researchers **(Figure 7A-C)**. Automated input processing sanitizes raw input strings, strips FASTA headers, filters non-canonical characters, and formats sequences with structural mask tokens (aa#) for tokenization. Two-stage ensemble inference executes the pipeline across the five-seed ensemble in under two seconds per sequence on a standard GPU, returning calibrated point estimates, aleatoric uncertainty, epistemic disagreement, and 95% confidence intervals for both T_m_ and OGT **(Figure 7B)**. Biophysical sequence profiling integrates a per-residue secondary structure propensity classifier that identifies α-helices, *β*-sheets, and contiguous flexible loop regions, highlighting potential initiation sites for thermal denaturation **(Figure 7C)**. Interactive visualization and export displays dynamic stability regime gauge charts, confidence interval bars, and provides downloadable CSV and JSON reports for high-throughput candidate libraries **(Figure 7C)**.

## Discussion

Predicting the thermal stability and environmental adaptability of proteins directly from sequence space remains a central objective in computational biology and protein engineering. While the emergence of self-supervised protein language models (pLMs) has significantly advanced predictive performance, relatively few studies have systematically tackled the interconnected biophysical, architectural, and evaluation bottlenecks in this domain. To our knowledge, current literature leaves several critical challenges largely unaddressed: the reliance on 1D sequence representations that fail to capture the 3D tertiary packing and flexible surface loops governing thermal denaturation, the destructive gradient interference that arises when jointly optimizing macroscopic host OGT and microscopic single-molecule T_m_ on a shared backbone, the severe natural data skew (>87% mesophiles) that drives networks into mesophilic prediction collapse, the over-reliance on whole-dataset error metrics that mask extremophile failure and conceal the necessity of bin-wise evaluation, and the absence of statistically calibrated uncertainty bounds required for reliable wet-lab decision-making. StableProt directly resolves these interconnected challenges by combining Foldseek 3Di structure-aware representations with parameter-isolated neural pathways, targeted 14% mesophilic subsampling, bin-resolved probabilistic scoring (CRPS), and observation-level uncertainty calibration.

A fundamental prerequisite for accurate stability prediction is accounting for the three-dimensional cooperativity of thermal denaturation. Unlike enzymatic function or family classification, which often correlate strongly with linear sequence motifs, thermal unfolding is governed by tertiary spatial phenomena such as core hydrophobic packing density, buried salt-bridge networks, backbone hydrogen bonding, and surface loop flexibility. While 1D sequence models achieve low error on in-domain test sets by memorizing linear sequence patterns, their predictive capacity degrades when transferring to out-of-distribution scaffolds. By mapping continuous 3D atomic coordinates into the discrete 20-state Foldseek structural alphabet, StableProt captures non-local packing contacts and backbone conformational strains that 1D sequence representations cannot resolve. In natural globins and industrial hydrolases, thermal unfolding typically initiates at flexible, solvent-exposed surface loops rather than within rigid secondary-structure cores. While 1D sequence encoders treat residues as linear strings, unbroken 3Di token sequences preserve these critical biophysical signals, enabling StableProt to resolve non-local initiation sites for thermal denaturation.

Beyond structural representation, our results demonstrate that predicting both T_m_ and OGT requires decoupling them at the architectural level. In earlier literature, OGT has frequently been treated as a direct proxy or co-equal multi-task target for T_m_ due to the vast abundance of microbiological growth records (∼1.4 million) relative to sparse biophysical unfolding measurements (<35,000). This pairing is conceptually complementary: OGT’s vast environmental abundance offsets Tm’s proteome-level scarcity, while Tm’s single-molecule precision resolves OGT’s species-level coarseness where all intra-cellular proteins share an identical host label. However, OGT and T_m_ represent distinct physical domains governed by disparate evolutionary pressures.OGT reflects a macroscopic ecological boundary shaped by membrane fluidity, metabolic cascades, and genomic GC content, whereas Tm represents a microscopic single-molecule denaturation endpoint. Consequently, mesophiles routinely express enzymes stable 30-50°C above host growth temperatures. Our gradient inner-product tracking demonstrates that forcing a shared neural trunk to optimize both targets simultaneously induces negative gradient correlation, creating destructive parameter competition. Crucially, by passing the predicted OGT scalar as an auxiliary prior link into the downstream T_m_ head at inference, the model allows macro-environmental context to inform single-molecule predictions without backward gradient corruption.

Beyond architectural modularity, StableProt demonstrates that targeted distribution-aware curation can outperform uncurated foundation scaling. Natural growth temperature databases are heavily skewed toward mesophiles, with over 87% of annotated organisms clustering between 25°C and 40°C. Under standard training regimes, neural networks minimize global loss by collapsing predictions toward this dense mesophilic prior mean. Competing predictors such as PRIME and ThermoFormer obtain deceptively low whole-dataset MAE by interpolating this mesophilic mode, yet their prediction errors surge roughly fourfold at elevated temperatures a frequency-dependent error pattern that recurred identically across three independent partitions: the internal holdout, decontaminated BacDive, and external zero-shot BRENDA. Because whole-dataset metrics are dominated by the mesophilic majority, this failure is completely masked until models are evaluated across 10°C temperature intervals (bin-wise MAE). Under this bin-balanced evaluation, StableProt outperforms competing models across seven of ten thermal intervals on BacDive (and six of ten on the full holdout), while maintaining a flat 0.76× collapse ratio on BRENDA. By subsampling mesophiles to 14% while retaining all extremophiles, StableProt balances batch gradients across thermal regimes, accelerating convergence and eliminating mesophilic collapse. Consequently, while models such as ThermoFormer require pre-training across 96 million uncurated sequences, StableProt achieves superior extremophile recall using fewer than 1 million curated records.

From a practical enzyme engineering standpoint, deterministic point predictions without uncertainty estimates provide inadequate guidance for wet-lab validation. A prediction of 65°C ± 2°C justifies immediate synthesis, whereas the same prediction with an uncertainty interval of ±18°C signals an out-of-distribution extrapolation that warrants caution. By coupling a heteroscedastic Huber-bounded Gaussian likelihood objective with a 5-seed deep ensemble, StableProt explicitly separates aleatoric data noise from epistemic model disagreement. Furthermore, calculating the true observation-level predictive variance resolves a widespread statistical flaw in existing literature: reporting the standard error of the ensemble mean artificially compresses intervals by √5 ≈ 2.24×, yielding nominal 95% confidence intervals that fail to cover more than ∼50% of real laboratory measurements. By preserving observation-level dispersion and applying cross-fitted scaling (c* = 1.65), StableProt delivers statistically reliable bounds. Across prospective evaluations spanning diverse industrial enzymes and environmental proteins, StableProt demonstrated this reliability by actively widening its confidence bounds for unannotated variants, successfully preventing false precision and ensuring laboratory resources are committed only to high-certainty candidates.

To ensure reliable interpretation, several biophysical boundary conditions should be acknowledged and are the subject of ongoing work. First, empirical Tm values vary with assay methodology (DSF, CD, DSC, TPP), heating rates, protein concentration, pH, and buffer ionic strength. Inverse-IQR weighting down-weights the most variable records, and StableProt accordingly reports intrinsic fold stability rather than condition-conditioned melting behavior; explicit pH- and buffer-aware multimodal modeling is a natural future direction. Second, oligomerization, protein-protein interfaces, and bound cofactors contribute substantially to stability in many systems, and the current implementation tokenizes monomeric ESMFold structures. Extending the structural token stream to multi-chain assemblies, for example via multimer-aware structure prediction, is planned future work for obligate oligomeric complexes. Third, on the FireProtDB single-point mutation benchmark (N = 3,649), the model achieved a directional accuracy of 56.8% (AUC = 0.585). This reflects an inherent boundary: StableProt is optimized for macroscopic candidate prioritization across sequence space, not for atomic-resolution enthalpy-entropy compensation. It is therefore positioned as an upstream candidate filter intended to be paired with downstream atomic-resolution refinement methods. Fourth, true psychrophilic enzymes (<25°C) remain severely underrepresented across public repositories (<2% of curated records), and cold-adaptation signatures remain a community data need; we anticipate that expanded psychrophile-focused metagenomic curation will improve coverage of this regime. Fifth, taxonomic coverage in the OGT training corpus remains biased toward cultivated bacteria and under-samples archaeal and uncultured extremophile lineages. Finally, the prospective enzyme suite reported here was assembled from curated experimental literature and publicly available biocatalyst panels, and *de novo* wet-lab characterization of novel Loop Engineering Toolkit designs by automated cell-free expression and high-throughput CD/DSF assays is designated as a second-generation experimental follow-up. To provide researchers with immediate access to these models, the entire dual-stage pipeline is hosted as an interactive web server (https://stableprot.dixitlab.org), delivering automated 3Di tokenization, two-stage ensemble inference, per-residue loop secondary structure profiling, and calibrated predictive interval.

In conclusion, StableProt resolves longstanding vulnerabilities in computational stability prediction, overcoming sequence representation limits, multi-task gradient interference, mesophilic collapse, and inappropriate global metrics. Engineered directly for protein engineers, enzymologists, synthetic biologists, and bioprospectors navigating uncharacterized sequence spaces, the platform minimizes experimental attrition prior to wet-lab synthesis by providing observation-level confidence bounds validated on prospective industrial enzymes. Freely accessible to the global community, the StableProt web server transforms stability screening into an actionable, risk-aware discovery pipeline. **Together, these results position StableProt as a statistically calibrated, broadly accurate, and robust predictor of protein thermal stability. It may help in the screening of thermostable enzymes from genomic databases, and engineering of novel biocatalysts.**

## Acknowledgements

The authors acknowledge funding support from DBT grant no BT/PR51568/BID/7/1044/2023. BRIC-ILS high performance computing facility is also acknowledged.

## Author contributions

Bibhu Prasad Behera and Anshuman Dixit conceptualized, designed the experiments and wrote the manuscript.

## Supplementary data

Supplementary data is available at **Supplementary Data.docx** file

## Conflict of interest

No conflict of interest

## Data availability

To ensure full reproducibility of all empirical benchmark comparisons while safeguarding proprietary pre-training data curation pipelines, our code, pre-trained models, and evaluation splits are distributed across the following open-source repositories;

(1) **GitHub Repository (Source Code & Inference Scripts)**: https://github.com/Bibhuprasadbehera/StableProt
(2) **Pre-Trained Model Weights (HuggingFace Hub)**: https://huggingface.co/bibhu111/StableProt
(3) **Evaluation Holdouts & Benchmark Splits (Zenodo DOI)**: https://zenodo.org/records/23916715

## Notes

https://stableprot.dixitlab.org/

### Competing Interest Statement

The authors have declared no competing interest.

