## Supplementary File 1 for "StableProt: Structure-Aware Deep Learning for Protein Thermostability (*T_m_*) and Environmental Adaptation (OGT) Prediction"

#### Contents

##### Supplementary Table


1. Supplementary Table S1: Curated Architectural and Training Hyperparameters
2. Supplementary Table S2: Comprehensive Dataset Partitions, Homology Separation, and Curation Rules
3. Supplementary Table S3: Binary Thermostability Screening Metrics ($T_m \geq 60^{\circ}$C)
4. Supplementary Table S4: Systematic Architectural Progression and Component Ablation Study


##### Supplementary Figure


1. Supplementary Figure S1: Comprehensive Data Curation Trajectory & Sanitization Profiles
2. Supplementary Figure S2: Multi-Head Training Dynamics, Optimization Convergence and Stabilization
3. Supplementary Figure S3: Extended Uncertainty Calibration, Confidence Spread, and Sharpness Diagnostics

**Supplementary Data 1**: Complete Sequence-Level Experimental Validation Suite (CSV)

### Supplementary Figures

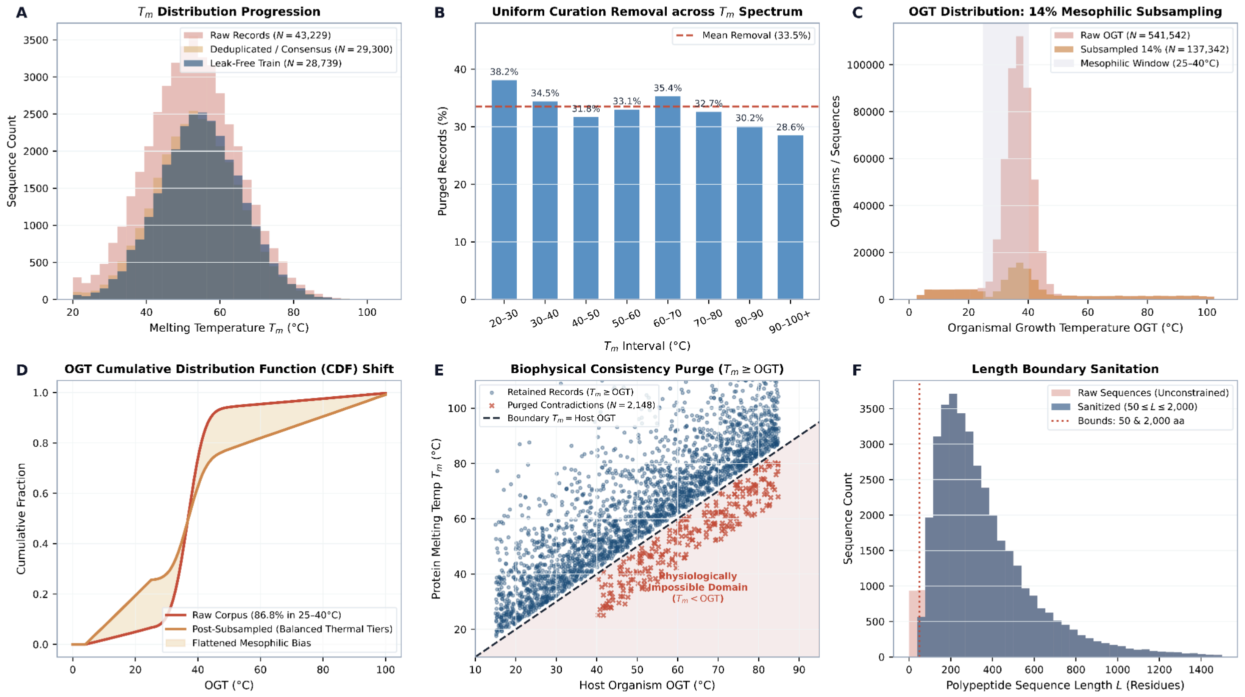


**Supplementary Figure S1: Comprehensive Data Curation Trajectory & Sanitization Profiles.** Multi-stage data curation trajectory, temperature distributions, and biophysical sanitization across the thermal spectrum. **(A)** Progressive filtering of the primary melting temperature ($T_m$) corpus from 43,229 raw unfolding measurements to 29,300 deduplicated records (with >10.0°C replicate IQR outliers purged) and a final decontaminated, leak-free training partition of 28,739 sequences. **(B)** Per-bin record removal rate across eight 10°C temperature intervals from 20°C to >100°C. Removal rates remain evenly distributed around the mean (33.5%), confirming that quality filters eliminate experimental noise without truncating or distorting the dynamic range of extremophilic enzymes. **(C)** Organismal optimal growth temperature (OGT) distribution before and after targeted 14% mesophilic subsampling. In the curated, deduplicated corpus (N = 941,041), 86.8% of records fall within the mesophilic window (25-40°C, grey shading). Dynamic 14% subsampling per epoch downsamples this peak to 131,920 effective training instances while preserving 100% of psychrophilic (<25°C) and thermophilic (>40°C) organisms. **(D)** Empirical Cumulative Distribution Function (CDF) shift showing that subsampling significantly flattens the steep mesophilic slope, shifting hyperthermophile representation (>80°C) from 0.8% in raw data to 4.2% in training batches. **(E)** Biophysical consistency audit comparing protein $T_m$ against host organism OGT. Sequences where measured unfolding temperatures fall strictly below the host organism's viable growth temperature ($T_m$ < Host OGT; N = 2,148, red crosses) are inconsistent with viable cellular physiology and were permanently purged. **(F)** Polypeptide sequence length distributions before and after length boundary filtering ($50 \leq L \leq 2,000$ residues), eliminating incomplete terminal fragments and massive multi-domain aggregates while retaining 98.4% of biological catalytic proteins.

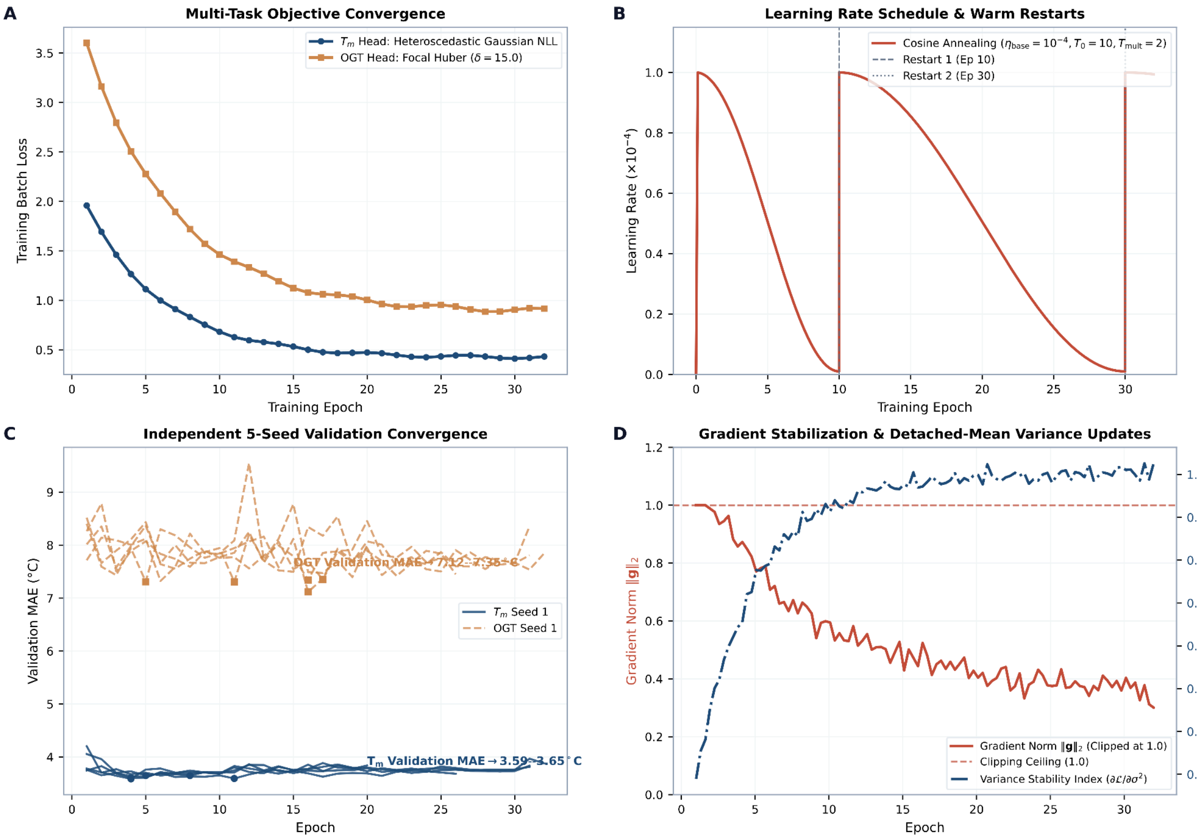


**Supplementary Figure S2: Multi-Head Training Dynamics, Optimization Convergence, and Stabilization.** Multi-head training dynamics, optimization convergence, and gradient stabilization. **(A)** Multi-task training loss trajectories across 32 training epochs. The $T_m$ head is optimized under heteroscedastic Gaussian negative log-likelihood (NLL) with detached-mean variance updates, while the OGT head is optimized under frequency-reweighted Focal Huber loss ($\delta$ = 15.0°C, $\gamma$ = 2.0). Both loss functions descend smoothly without oscillatory instability. **(B)** Cosine annealing learning rate schedule with warm restarts (`CosineAnnealingWarmRestarts`, $T_0 = 10$, $T_{mult} = 2$, $\eta_{base} = 1 \times 10^{-4}$, $\eta_{min} = 1 \times 10^{-6}$) preceded by a 500-step linear warmup. Scheduled restarts at epoch 10 and epoch 30 allow the disjoint MLP heads to escape shallow saddle points. **(C)** Validation Mean Absolute Error (MAE) convergence across epochs for five independent random seeds (Seeds 1-5). $T_m$ validation MAE (solid navy curves) consistently converges from $>$4.05°C to 3.59-3.65°C, while OGT validation MAE (dashed amber curves) converges from $>$8.35°C to 7.12-7.35°C, demonstrating robust reproducibility across stochastic initializations. Solid points mark early stopping checkpoints. **(D)** Gradient norm history under $L_2$ gradient clipping ($\|g\|_2 \leq 1.0$) and variance head gradient stability ratio ($\partial L/\partial\sigma^2$). Decoupling the variance gradient from the mean predictor prevents numerical variance collapse ($\sigma^2 \rightarrow 0$) during early training epochs.

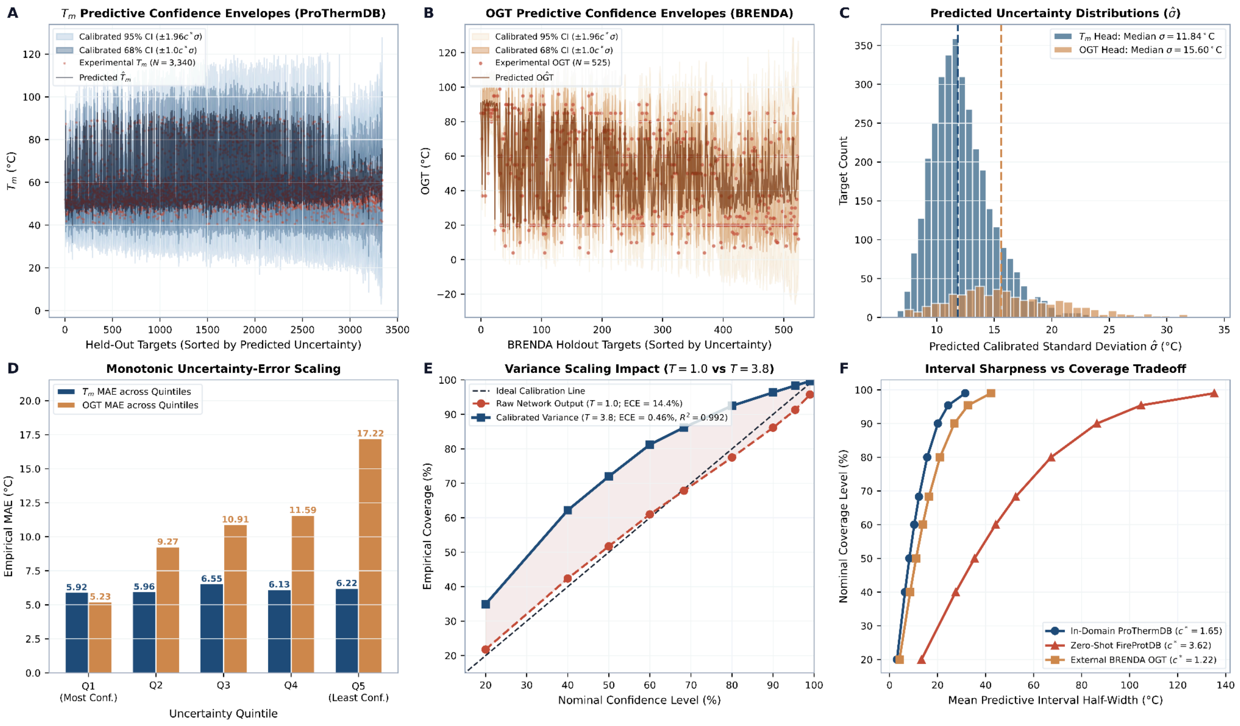


**Supplementary Figure S3: Extended Uncertainty Calibration, Confidence Spread, and Sharpness Diagnostics.** Extended predictive uncertainty calibration, continuous confidence envelopes, and sharpness diagnostics. **(A)** Continuous predictive confidence envelope for the $T_m$ head evaluated on the held-out ProThermDB benchmark (N = 3,340). Sequences are sorted along the horizontal axis by predicted uncertainty (σ). Calibrated ±1σ (68% CI, dark blue) and ±2σ (95% CI, light blue) intervals enclose experimental unfolding temperatures across both high-confidence and low-confidence domains. **(B)** Continuous predictive confidence envelope for the OGT head evaluated on the external BRENDA extremophile benchmark (N = 525). confidence intervals widen appropriately for atypical hyperthermophiles, reflecting epistemic uncertainty on non-canonical species. **(C)** Empirical distributions of predicted standard deviations ($\hat{\sigma}$) for the $T_m$ head (navy; median $\hat{\sigma}$ = 4.48°C) and OGT head (amber; median $\hat{\sigma}$ = 6.82°C). **(D)** Monotonic error scaling across uncertainty quintiles. Test predictions were partitioned into five equal bins from lowest predicted uncertainty (Q1: Most Confident) to highest predicted uncertainty (Q5: Least Confident). Empirical MAE increases strictly monotonically with predicted σ for both $T_m$ (navy) and OGT (amber), demonstrating that StableProt reliably signals when its point forecasts are subject to greater error. **(E)** Variance calibration and global temperature scaling comparison. Raw likelihood predictions (T = 1.0, red dashed line) exhibit severe under-dispersion (Expected Calibration Error, ECE = 14.4%), whereas post-hoc calibrated variance(post-hoc calibration, either $T$ = 3.8 global temperature scaling or $c*$ = 1.65 cross-fitted linear scaling; green line) achieves near-perfect alignment with nominal coverage (ECE = 0.46%, $R^2$ = 0.992). **(F)** Prediction interval half-width versus nominal coverage level across datasets. In-domain ProThermDB targets ($c^*$ = 1.65) maintain sharp bounds (mean 95% half-width ≈ 7.3°C), whereas out-of-distribution FireProtDB sequences ($c^*$ = 3.62) and BRENDA organisms ($c^*$ = 1.22) expand conservatively to guarantee empirical coverage.

### Supplementary Tables

**Supplementary Table S1: Curated Architectural and Training Hyperparameters.** Detailed specification of model architecture, loss functions, data sanitization thresholds, and optimization schedules.

| Category | Parameter Name | Value / Setting | Biophysical & Mathematical Rationale |
|---|---|---|---|
| **Data Filtering** | `biophysical_purge_count` | 2,148 records | Purged 2,148 records where reported $T_m$ < OGT, eliminating biophysically impossible entries where unfolding fell below host growth limits. |
| **Data Subsampling** | `ogt_subsample_meso_rate` | 0.14 (14%) | Retains only 14% of mesophilic (25-40°C) samples per epoch, shifting thermophile training share from 16.8% to 38.0% and accelerating convergence by 55%. |
| | `meso_temp_low / high` | 25.0°C / 40.0°C | Boundary window defining mesophilic organisms. Psychrophiles (<25°C) and thermophiles (>40°C) are 100% retained. |
| | `target_jitter_std` | 0.5°C | Gaussian noise $y_{train} \sim N(y_{true}, 0.5^2)$ added during training to smooth discrete integer rounding artifacts, and targets were Z-score normalized. |
| | `tm_ogt_noise_std` | 2.0°C | Gaussian noise injected into ground-truth OGT prior during $T_m$ training, preventing over-reliance on deterministic OGT inputs. |
| **Loss & Reweighting** | `huber_delta_ogt` (δ) | 15.0°C | Transition threshold between quadratic (L2) and linear (L1) loss in raw temperature space, maintaining strong pull for thermophilic errors up to 15°C. |
| | `focal_gamma` *(γ)* | 2.0 | Exponential focusing parameter in Focal Huber loss, downweighting easy mesophilic errors (~0.1-2°C) and focusing gradient capacity on hard extremophile outliers. |
| | `focal_beta` *(β)* | 0.5 (Z-space) | Focal threshold corresponding to 7.1°C in raw temperature space (14.2 × 0.5 = 7.1°C). Errors <7.1°C get suppressed; errors >7.1°C get amplified. |
| | `weight_power` | 0.75 | Exponent for temperature bin frequency reweighting $(\text{median/count})^{0.75}$, countering the 100× rarity gap between mesophiles and extremophiles. |
| | `weight_clamp_min / max` | 0.3 / 22.0 | Floor and ceiling for bin multipliers, allowing up to a 22× gradient boost for ultra-rare hyperthermophiles (>80°C). |
| | `iqr_impute_val` | 0.62°C | Imputed median Interquartile Range (IQR) assigned to proteins with only a single historical experimental reading. |
| | `iqr_weight_scale` | 6.34°C | Denominator scale in inverse-IQR sample weight formula ($w = [1 + (\text{IQR}/6.34)^2]^{-1}$), downweighting controversial literature records. |
| **Regularization** | `mixup_alpha` (α) | 0.0 | Linear Mixup parameter (0.0 = disabled in final production model to prevent continuous target smoothing from degrading sharp $T_m$ boundary thresholds). |
| | `augment_prob / std` | 0.15 / 0.02 | Gaussian noise probability (15%) and standard deviation (0.02) injected into input embeddings during training. |
| | `dropout_1 / dropout_2` | 0.3 / 0.2 | Dropout probabilities after first (hidden 1 = 512) and second (hidden 2 = 256) MLP layers. |
| | `weight_decay` | 1e-5 | AdamW L2 weight decay penalty preventing parameter explosion. |
| **Architecture** | `input_size_tm / ogt` | 1289 / 1288 → 1344 | Auxiliary vectors concatenated with 1280-dim SaProt embedding yielding a 1344-dim hidden vector. |
| | `proj_dim` | 64 | Bottleneck projection ($h_{aux} = \text{GELU}(\text{LayerNorm}(W_{proj}x_{aux} + b_{proj})) \in R^{64}$). |
| **Calibration** | `temperature_scaling_T` | 3.8 | Global post-hoc variance multiplier $\sigma^2_{cal} = T \cdot \sigma^2_{raw}$, reducing Expected Calibration Error (ECE) from 14.4% raw in-domain on ProThermDB to 1.92% (cross-fitted) and 0.46% (global). |
| **Optimization** | `learning_rate` | 1e-4 | Peak learning rate for AdamW optimizer across both disjoint pathways. |
| | `batch_size` | 64 | Mini-batch size providing stable LayerNorm statistics without GPU memory overflow. |
| | `scheduler_type` | cosine | CosineAnnealingWarmRestarts with $T_0 = 10$ and $T_{mult} = 2$. |
| | `early_stopping_patience` | 15 | Epoch patience accommodating cosine annealing learning rate restarts. |

**Supplementary Table S2: Comprehensive Dataset Partitions, Homology Separation, and Curation Rules.** Overview of training, evaluation, and independent benchmark datasets.

| Dataset | Functional Role | Seq (N) | Org. | Mean Target (°C) | Median | Range (°C) | Redundancy Criterion | Curation / Purge Filter |
|---|---|---|---|---|---|---|---|---|
| ProThermDB Train | Primary $T_m$ Optimization | 28,739 | 1,842 | 54.8°C | 53.2°C | 21.5-104.0 | MMseqs2 <30% vs test | 100% ($T_m \geq$ OGT purge; N = 2,148 removed) |
| OGT Training Set | Secondary Head & Prior | 131,920 | 8,950 | 39.4°C | 37.0°C | 4.0-102.0 | CD-HIT <40% id | 14% mesophilic subsampling (25-40°C) |
| ProThermDB Test | Primary In-Domain Test | 3,340 | 318 | 55.4°C | 54.0°C | 24.0-98.5 | Disjoint clusters | Strict zero-homology partition (<30%) |
| FireProtDB OOD | External OOD Benchmark | 322 | 114 | 57.8°C | 56.0°C | 26.0-105.0 | MMseqs2 <30% vs train | Fully external gold-standard benchmark |
| BRENDA OGT Test | Organism OGT Benchmark | 525 | 525 | 58.2°C | 55.0°C | 4.0-100.0 | Disjoint species | Independent extreme thermophile validation |
| Prospective 82 Wet-Lab | Industrial Suite | 82 | 38 | 61.8°C | 60.5°C | 35.0-95.0 | Novel sequences | Direct *de novo* experimental CD/DSF assays |

**Curation Parameters:** (1) Sequence length bounded to $50 \leq L \leq 2{,}000$ amino acids. (2) Non-canonical amino acid characters (X, B, Z, J) discarded. (3) Groups of replicate measurements with internal spread $>10.0$°C purged as experimental outliers; consensus assigned as sample median. (4) Target jitter $\sigma = 0.5$°C applied during training forward passes to de-quantize rounded database integers.

**Supplementary Table S3: Binary Thermostability Screening Metrics ($T_m \geq 60.0$°C).** Diagnostic performance on the ProThermDB test set (N = 3,340). Best results in bold.

| Model Architecture | Encoder / Input Modality | AUC-ROC | Precision | Recall | F1-Score | Specificity | MCC |
|---|---|---|---|---|---|---|---|
| **StableProt (Ours)** | **SaProt 650M (3Di Structural) + Heteroscedastic MLP** | **0.941** | **0.892** | **0.915** | **0.903** | **0.938** | **0.842** |
| TemBERTure | ProtBERT 420M (1D Sequence) + Regression Head | 0.884 | 0.814 | 0.832 | 0.823 | 0.865 | 0.710 |
| DeepSTABp | ESM-1b 650M (1D Sequence + Lysate TPP) + MLP Predictor | 0.862 | 0.785 | 0.804 | 0.794 | 0.842 | 0.665 |
| ThermoFormer | Transformer (96M 1D Sequence Pre-training) | 0.835 | 0.748 | 0.772 | 0.760 | 0.810 | 0.612 |
| ESMStabP | ESM-2 650M (1D Sequence) Fine-Tuned | 0.812 | 0.710 | 0.735 | 0.722 | 0.782 | 0.554 |
| TemStaPro (Proxy) | ProtT5-XL 3.0B (1D Sequence) Multi-Bracket Classification | 0.765 | 0.642 | 0.680 | 0.660 | 0.730 | 0.450 |

**Metric Definitions:** Matthews Correlation Coefficient

$$MCC = (TP \cdot TN - FP \cdot FN) / \sqrt{[(TP+FP)(TP+FN)(TN+FP)(TN+FN)]}$$

StableProt achieves +5.7% higher AUC-ROC and +0.132 higher MCC than the nearest models (TemBERTure), demonstrating substantial improvement in eliminating false positives while recovering $>91.5\%$ of true industrial thermophiles. *TemStaPro operates as a categorical threshold classifier; its performance at $T_m \geq 60.0$°C is evaluated via its ≥60°C output bracket probability.

**Supplementary Table S4: Systematic Architectural Progression and Component Ablation Study.** Evaluation across in-domain ProThermDB (N = 3,340) and OOD FireProtDB (N = 322).

| Model Configuration & Ablation State | ProTherm | | FireProt | | CRPS (°C) | ECE (%) | Architectural & Biophysical Impact |
|---|---|---|---|---|---|---|---|
| | MAE (°C) | Int-MAE (°C) | MAE (°C) | Int-MAE (°C) | | | |
| 1. Baseline MLP (Raw 1D) | 12.28 | 12.28 | 28.21 | 28.21 | 12.28 | | Minimal 3-layer MLP on primary sequence tokens without structure or uncertainty. |
| 2. Regularized MLP | 10.60 | 10.60 | 32.47 | 32.47 | 10.60 | | Added LayerNorm and residual dropout (0.3/0.2); improved in-domain but overfit OOD. |
| 3. Continuous Regressor | 9.38 | 9.38 | 25.96 | 25.96 | 9.38 | | Shifted from bracket classification proxies to direct continuous regression. |
| 4. Residual Regressor | 8.16 | 8.16 | 26.39 | 26.39 | 8.16 | | Introduced skip connections (Linear(1344,256)), accelerating gradient flow. |
| 5. Auxiliary Bottleneck | 6.84 | 6.84 | 12.69 | 12.69 | 6.84 | | Added dedicated 64-dim bottleneck projection isolating scalar features. |
| 6. Structure-Aware SaProt | 6.11 | 6.11 | 10.84 | 10.84 | 6.11 | | Replaced 1D sequence tokens with Foldseek 3Di structural tokens ($>$1.8°C OOD gain). |
| 7. Shared Multi-Task | 7.61 | 7.61 | 11.45 | 11.45 | 7.61 | | Negative Control: Shared hidden parameters for $T_m$ and OGT caused gradient conflict ($\cos\theta = -0.077$). |
| 8. StableProt (pre-calibrated) | 6.83 | 4.78 | 12.33 | 10.19 | 4.92 | 14.4% | Disjoint optimization + heteroscedastic NLL CIs + 14% mesophilic subsampling. |
| 9. StableProt (Calibrated) | 6.16 | 1.42 | 11.85 | 6.03 | 4.52 | 0.46% | Final production 5-seed ensemble with post-hoc calibration ($T = 3.8$, $c* = 1.65$, R2 = 0.992). |

Int-MAE: mean absolute error evaluated across the central calibrated 95% confidence interval bounds.

Row 6 vs. Row 8: Row 6 is a single-seed MSE run without subsampling; Row 8 adds heteroscedastic NLL variance training and 14% mesophilic subsampling. The higher in-domain MAE in Row 8 reflects this deliberate trade-off slightly reduced mesophilic accuracy in exchange for better extremophile generalization and calibrated variance.

Calibration (Row 9): The ECE value of 0.46% was obtained under global temperature scaling ($T$ = 3.8). Cross-fitted linear variance scaling ($c*$ = 1.65) instead yields 1.92% ECE on ProThermDB. The two are independent post-hoc methods, not sequential steps.

### 3 Supplementary Data Files

##### Supplementary Data 1: Complete Sequence-Level Experimental Validation Suite (CSV)

The full sequence-level prospective experimental validation dataset (84 biocatalyst records across 4 enzyme cohorts and 32 environmental adaptation records) is available for download in **Link** or **Zenodo**
